# Genome-wide CRISPRi-seq maps essential-gene contributions to aminoglycoside susceptibility in *Pseudomonas aeruginosa*

**DOI:** 10.64898/2026.11.01.763562

**Authors:** Weronika Czekała, Jan Gawor, Raymi Edgar Goitia Camacho, Barbara Domańska, Luis Felipe Padilla Martinez, Karolina Żuchniewicz, Filip Pośnik, Aneta Agnieszka Bartosik, Xue Liu, Adam Kawałek

**Affiliations:** Institute of Biochemistry and Biophysics, Polish Academy of Sciences, Warsaw, Poland; Doctoral School of Molecular Biology and Biological Chemistry at IBB PAS, Warsaw, Poland; Faculty of Chemistry, Chair of Drug and Cosmetics Biotechnology, Warsaw University of Technology, Poland; Guangdong Provincial Key Laboratory of Regional Immunity and Diseases, Department of Pathogen Biology, Shenzhen University Medical School, Shenzhen, Guangdong, China

## Abstract

*Pseudomonas aeruginosa* is an opportunistic ESKAPE pathogen whose multiple intrinsic and acquired antibiotic resistance mechanisms restrict treatment options. Essential genes offer potential antimicrobial targets, but their vulnerability to partial inhibition and contribution to antibiotic susceptibility remain incompletely characterized. Here, we developed a rhamnose-inducible CRISPR interference (CRISPRi) platform and an 83,078-guide library for genome-wide screening in PAO1, with mismatch guides enabling graded repression. Fitness profiling in antibiotic-free Mueller–Hinton medium identified 743 genes whose repression reduced competitive fitness and revealed sequence features associated with guide performance. Combining full-match and mismatch-guide responses revealed substantial variation in essential-gene fitness effects across predicted repression strengths, while comparison of fitness profiles between Mueller–Hinton medium and fetal bovine serum identified condition-dependent effects alongside a conserved fitness landscape.

Under aminoglycoside exposure, we identified 118 recurrent sensitizing and 120 recurrent protective gene knockdowns, including many targeting essential and fitness-associated functions. Growth and killing assays with individual strains confirmed that CRISPRi targeting of *atpI*, *accC*, *secA* and *ftsZ* enhanced tobramycin activity. Finally, transcriptome analysis of these knockdowns confirmed target silencing, captured polar effects on neighboring genes and revealed distinct but overlapping transcriptional responses. Together, these findings quantify the variable fitness costs of essential-gene repression and identify candidate targets for potentiating aminoglycoside activity.

**IMPORTANCE:** Essential bacterial genes are difficult to study because their complete disruption prevents growth. Yet knowing that a gene is essential does not reveal how strongly its activity must be reduced to impair growth. Here, we developed a method to examine how reduced gene expression affects fitness across the genome of *Pseudomonas aeruginosa*, a major cause of difficult-to-treat infections. This approach identified essential functions particularly vulnerable to partial depletion and mapped how gene repression alters aminoglycoside susceptibility. Repressing selected genes impaired growth and disproportionately increased killing by tobramycin. These tools and datasets provide a resource for prioritizing candidate antibacterial targets and investigating whether partial inhibition of essential functions can enhance the activity of existing antibiotics.

## INTRODUCTION

Many antibacterial agents act by inhibiting essential cellular processes, therefore the essential gene products represent candidate antibacterial targets for novel approaches (1–3). Genome-wide approaches, particularly transposon-insertion sequencing (Tn-seq), identify genes required under defined growth conditions, while comparative analyses assess the conservation of essential functions across organisms (4, 5). Together, these approaches distinguish broadly conserved core essential genes from conditionally essential genes, required only in a particular environmental or genetic context (6). However, essentiality is generally assigned as a binary variable, with genes classified as either essential or non-essential (6). Because transposon-insertion screens primarily assess loss-of-function phenotypes, they do not directly predict the consequences of partial target suppression. Importantly, essential targets differ in how much depletion they tolerate before fitness declines (7–9). This quantitative response, termed vulnerability, can therefore guide target prioritization beyond binary essentiality classifications.

*Pseudomonas aeruginosa* is a versatile Gram-negative opportunistic pathogen and a member of the ESKAPE group of major healthcare-associated pathogens (10, 11). *P. aeruginosa* causes difficult-to-treat acute and chronic infections, particularly in hospitalized or immunocompromised individuals and in people with cystic fibrosis (12). Treatment is complicated by intrinsic resistance to multiple antibiotic classes and by the capacity to acquire additional resistance, emphasizing the need for new antibacterial targets or treatment combinations (10, 12). Essential and conditionally essential functions in *P*. *aeruginosa* have been identified through transposon-based analyses performed in laboratory media and infection-relevant environments (13–15). Additional mutant-library and Tn-seq screens have defined genes required during infection, biofilm growth, nutrient limitation and antibiotic exposure (16–25). Notably, the Tn-seq analysis across nine strains and five media identified 321 core essential genes, providing a conserved reference set of candidate antibacterial targets (14). Although essential genes can modulate fitness during antibiotic exposure, how their graded repression shapes antibiotic responses in *P. aeruginosa* remains poorly understood.

CRISPR interference (CRISPRi) provides an approach for analysing the fitness consequences of essential-gene repression and identifying genes and pathways particularly sensitive to inhibition. In CRISPRi, a single-guide RNA (sgRNA) containing a target-complementary spacer directs catalytically inactive nuclease Cas9 (dCas9) to a genomic sequence located next to a protospacer adjacent motif (PAM). Binding within promoter or regulatory regions can interfere with transcription initiation, whereas binding within transcribed regions can sterically block transcription elongation (26–28). For effective repression within coding regions using *Streptococcus pyogenes* dCas9 (dCas9_Spy_), 20-nucleotide spacers are typically designed to target the non-template strand (27).

Because targeting is specified by the sgRNA spacer, CRISPRi can be readily adapted to pooled or arrayed genome-scale libraries (26, 29–32). In pooled screens, sgRNAs serve as molecular barcodes whose relative abundance can be quantified by high-throughput sequencing, enabling parallel measurement of knockdown-associated fitness effects across conditions. Genome-scale CRISPRi screens have quantified fitness effects, essential-gene vulnerability and gene–drug interactions in diverse bacteria (26, 8, 29, 9, 33–38). The repression strength can be tuned by controlling dCas9 or sgRNA expression, introducing spacer mismatches, modifying the sgRNA scaffold or, for some dCas9 variants, selecting target sites with different PAM sequences (8, 39–42). This tunability enables graded analysis of essential-gene phenotypes and their effects on fitness and antibiotic susceptibility.

CRISPRi has previously been implemented in *P. aeruginosa* (43, 44). Recent studies have enabled genome-wide CRISPRi using different dCas9 variants and induction strategies. Zhang and co-authors developed a doxycycline-activated dCas9_Spy_ system in PA14 and applied it to identify determinants of gallium susceptibility (45). Kaczmarczyk and co-workers developed a CRISPRi system using IPTG-controlled *Streptococcus pasteurianus* dCas9 and constructed a library of more than 80,000 sgRNAs, which was subsequently used to identify genes involved in resistance to type VI secretion system attacks (46, 47). Sullivan and co-authors independently optimized a *S*. *pasteurianus* dCas9 system in which dCas9 and sgRNA expression were separately controlled by rhamnose-inducible promoters, enabling regulated depletion of essential genes (48). These studies establish CRISPRi for genome-wide functional analysis in *P. aeruginosa*. However, how graded repression of essential functions shapes antibiotic susceptibility remains incompletely defined.

Here, we describe a rhamnose-inducible dCas9_Spy_-based CRISPRi platform, and a genome-wide full-match and mismatch-guide library in PAO1. We used the library to quantify fitness effects across predicted repression strengths, identify sequence features associated with guide performance and map gene–aminoglycoside interactions, followed by validation of selected sensitizing knockdowns.

## MATERIALS AND METHODS

### Bacterial strains and culture conditions

Bacterial strains, plasmids and oligonucleotides used or constructed in this study are listed in Supplementary Table S1. Selected CRISPRi plasmids and the library will be deposited in Addgene. *Pseudomonas aeruginosa* PAO1 was used for pooled screening and most validation assays. Resequencing of the PAO1 clone used here revealed sequence differences relative to the PAO1-UW reference genome (NC_002516.2), including the ∼1-kb *PA4684*–*PA4685* (*mksEF*) deletion and a distinct *mexT* allele (Supplementary Table S1). Selected assays were also performed in PA14 and LESB58 obtained from BCCM (https://bccm.belspo.be/). *Escherichia coli* DH5α was used for routine cloning and plasmid propagation, and *E*. *coli* S17-1 λpir was used for conjugative transfer of suicide, and integrative plasmids into *P*. *aeruginosa*. Bacterial strains were routinely grown in L broth (LB; 1% tryptone, 0.5% yeast extract, 0.5% NaCl) at 37°C or in cation adjusted Mueller Hinton Broth 2 (Millipore, cat. 90922-500G). Growth assays were performed in microplate format using a Varioskan LUX multimode microplate reader (Thermo Scientific) by monitoring optical density at 600 nm (OD₆₀₀) over time at 37°C with orbital shaking. Unless indicated otherwise, overnight cultures in LB were diluted 1:200 into MH, supplemented with indicated concentrations of rhamnose and/or antibiotics. The serum-containing medium consisted of 60% fetal bovine serum (Value FBS, Gibco, Ref. A5209402) diluted in PBS (137 mM NaCl, 2.7 mM KCl, 10 mM Na₂HPO₄, and 1.8 mM KH₂PO₄, pH 7.4). Before shifting to serum-containing medium, cells were washed once with PBS. When required, antibiotics were used at the following final concentrations for *P. aeruginosa*: 100 μg/mL tetracycline, 300 μg/mL carbenicillin, and 30 μg/mL gentamicin for PAO1 / PA14 or 150 μg/mL gentamicin for LESB58. Triclosan 25 μg/mL was used for donor counterselection during bacterial conjugations. Expression from *araBAD*p-, *rha*p-, and *tac*p-based constructs was induced with L-arabinose, L-rhamnose, or IPTG, respectively.

### Genetic manipulations of bacteria

The PAO1 Δ*mexXY* mutant was constructed by markerless allelic exchange using the suicide vector pAKE600 essentially as described previously (49, 50). DNA fragments flanking *mexXY* were assembled in pAKE600 and plasmid was moved into PAO1 using *E. coli* S17-1 λpir. Integrants were selected with carbenicillin and triclosan. Double-crossover recombinants were obtained by 10% sucrose counterselection and screened by PCR.

The CRISPRi system developed here involved separate integration of dCas9_Spy_ and sgRNA cassettes. A mini-CTX derivative pAKW2268, containing codon-optimized *Streptococcus pyogenes dcas9* variant under control of the rhamnose-inducible *rha*p, was introduced into the chromosome using conjugation and integrants were selected with tetracycline. The vector backbone including tetracycline-resistance cassette was removed by Flp-mediated recombination with the use of plasmid pAK1555. Individual sgRNA cassettes were introduced separately using pAKW2049 or pAKW2235 mini-Tn7 vectors, derived from pUC18T-mini-Tn7T and pJMP1237 ((44, 51), Fig1A). Annealed spacer oligonucleotides were inserted by BsaI-mediated Golden Gate assembly, replacing the *lacZ*α or *mScarlet* screening cassette in pAKW2049 or pAKW2235, respectively. Exact plasmids, guide constructs and oligonucleotide sequences used in this study are listed in Supplementary Table S1.

sgRNA constructs were introduced into the corresponding dCas9-expressing *P. aeruginosa* strains by mini-Tn7-mediated chromosomal integration using triparental conjugation, based on established mini-Tn7 procedures (51, 52). An *E. coli* S17-1 λpir strain carrying the mini-Tn7 cargo vector and a S17-1 λpir helper strain carrying pJMP1039, encoding the TnsABCD transposition machinery, were used as donors (44). Following mating on LB agar, integrants were selected with gentamicin on LB containing triclosan for donor counterselection. For PAO1 and PA14, recipient cells were grown in liquid LB at 42°C before mating, whereas LESB58 was grown at 37°C.

### CRISPRi library construction and pooled screening

The PAO1 83k CRISPRi library was designed using candidate full-match sgRNAs generated with the Mobile-CRISPRi sgRNA-design workflow (44), CRISPRi-Pasteur (53), and the guide-design pipeline developed for mismatch-CRISPRi (8, 41). Genes were prioritized for increased guide coverage based on previously reported essentiality and fitness datasets (Supplementary Table S2, (13–16, 25, 54)). Guides supported by multiple design approaches, and with lower predicted off-target burden were selected preferentially. Candidate guides were restricted to the preferred CRISPRi strand orientation and the first 2.5 kb of coding regions, with mismatch-CRISPRi candidates additionally filtered to 25–70% GC content. Single-mismatch variants were chosen to span a range of predicted repression efficacies. Non-targeting controls were random 20-nt sequences with no PAO1 genomic matches of ≥11 nt detected using BLASTN with the blastn-short task. Library composition and guide sequences are provided in Supplementary Table S3. The final library design comprised 83,078 sgRNAs targeting 5,557 genes, including 26,602 full-match guides, 54,783 mismatch variants and 1,693 non-targeting controls. The synthesized oligonucleotide pool was cloned into pAKW2049 and transferred to the *rha*p-*dcas9*_Spy_ containing PAO1 strain by mini-Tn7-mediated integration. Approximately 2.5 × 10⁸ integrants were recovered, providing approximately 3,000-fold representation of the designed library. The pooled library was stored in LB with 16% glycerol at −80°C. Detailed cloning procedures are provided in Supplementary Methods.

For pooled screening, library aliquots were inoculated into MH at OD₆₀₀ ≈ 0.05 and pre-induced with 0.1% rhamnose for 2 h, reaching OD₆₀₀ ≈ 0.3. Samples collected after pre-induction were designated T0 and served as the reference for subsequent fitness comparisons. Cultures were then diluted 1:10 into the respective screening media containing 0.1% rhamnose. All screening cultures had a final volume of 20 mL. Untreated MH cultures and aminoglycoside-treated cultures were propagated for 16 h to generate T1, corresponding to approximately seven generations after T0 under untreated conditions. Aminoglycoside treatments comprised tobramycin at 0.025, 0.1, 0.4 or 1 µg mL⁻¹ (MIC 1 µg mL⁻¹) and amikacin at 2 µg mL⁻¹ (MIC 4 µg mL⁻¹), with each treatment compared with its matched induced untreated control. To generate T2 populations in B1, untreated and TOB-treated T1 cultures were diluted 1:128 into fresh MH containing 0.1% rhamnose, without or with 0.4 µg mL⁻¹ TOB, respectively, and propagated for a 20h, corresponding to approximately 14 generations after T0. To assess condition-dependent fitness, the library was transferred in parallel into MH or 60% FBS medium, with FBS cultures propagated for 24 h. Library screens were performed with two or three biological replicates per condition and batch.

### Phenotypic characterisation of CRISPRi strains

CRISPRi effect was assessed using strains carrying selected full-match and mismatched sgRNAs. Strains were grown in MH with or without 0.1% rhamnose, and OD₆₀₀ was monitored using a Varioskan LUX microplate reader. For reporter silencing analysis, strains were grown in LB. mRFP fluorescence (excitation: 580 nm; emission: 610 nm) was normalized to OD₆₀₀ and summarized over the 10–15 h analysis window. To validate tobramycin sensitization, strains were grown in MH under four conditions combining the presence or absence of 0.1% rhamnose and 0.4 μg mL⁻¹ tobramycin. Growth rates were calculated from blank-corrected growth curves over experiment-specific time windows. The expected combined-treatment growth rate was calculated by multiplying the mean growth rates under CRISPRi induction alone and TOB alone, then dividing by the mean untreated growth rate. Combined-treatment growth rates were compared with this expectation using a one-sided, one-sample *t*-test. Significant effect required *p* < 0.05 and a reduction below expectation of ≥10% of the untreated growth rate, since strong inhibition by CRISPRi alone can bring the expectation close to zero, limiting detection of additional tobramycin effects. For killing assays, overnight LB cultures were diluted 1:200 into MH containing 0.1% rhamnose, grown for 3 h at 37°C and subsequently diluted 1:10 into fresh MH containing 0.1% rhamnose with or without the indicated concentration of tobramycin. Viability was determined as CFU mL⁻¹ in three biological replicates. Treatment-associated reductions in viable counts were compared between CRISPRi knockdown and non-targeting control strains using paired, two-sided *t*-tests matched by biological replicate.

### CRISPRi-seq

Genomic DNA was extracted from library samples, and sgRNA spacer regions were amplified, indexed and sequenced as described in detail in Supplementary Methods. Data analysis was adapted from Ward et al. (34). Reads containing the sgRNA cassette were selected using Cutadapt (55), and guide abundances were quantified by exact 20-nt matching with Seal.sh from BBTools (56). Guide-level differential abundance was analysed with edgeR (57), and unadjusted guide-level *p* values were combined at the gene level using Stouffer’s method implemented in poolr (58). Detailed description of data analysis flow is included in Supplementary Methods. Full-match guide counts were also analysed using MAGeCK RRA (59), with median normalization and gene-level effects calculated as the median log₂ fold change across guides targeting each gene. Raw CRISPRi-seq sequencing data generated in this study are available through the NCBI Gene Expression Omnibus (GEO) under accession GSE349013 (accessed 1.10.2026, https://www.ncbi.nlm.nih.gov/geo/query/acc.cgi?acc=GSE349013).

### Outer-membrane permeability assay

Outer-membrane permeability was assessed using 1-N-phenylnaphthylamine (NPN), adapting the method of Sposato et al., (60). Overnight cultures of PAO1 CRISPRi strains and the non-targeting control were diluted 1:200 in MH containing 0.1% rhamnose and incubated at 37°C for 3 h. Cells were harvested by centrifugation, washed and resuspended in 5 mM HEPES (pH 7.2), and normalized to OD₆₀₀ = 0.2. In microplate wells, 100 µL of cell suspension was mixed with 100 µL of buffer containing 20 µM NPN (10 µM final concentration) or NPN-free buffer as an autofluorescence control. After 2 min, fluorescence was measured at excitation/emission wavelengths of 350/420 nm using Varioskan LUX microplate reader. Fluorescence was normalized to OD₆₀₀ and expressed relative to the corresponding non-targeting control.

### RNA-sequencing

For RNA-seq analysis, PAO1 CRISPRi strains carrying *atpI*_g1m2, *accC*_g1m3, *secA*_g1m3, *ftsZ*_g1m4, or the non-targeting control_1 guide were grown overnight in MH medium at 37°C with shaking. Overnight cultures were diluted 1:200 into fresh MH supplemented with 0.1% rhamnose and grown for 3 h, reaching an OD₆₀₀ ≈ 0.3. In parallel, the control_1 harbouring strain was grown under the same conditions in MH containing 0.1% rhamnose and 0.4 µg mL⁻¹ tobramycin (TOB) to determine the transcriptional response to TOB alone. For each condition, total RNA was isolated from 2 mL of culture from three independent biological replicates. Cells were stabilized with RNAprotect Bacteria Reagent (Qiagen), and total RNA was purified using the RNeasy Mini Kit (Qiagen). Residual genomic DNA was removed using the TURBO DNA-free Kit (Ambion), and DNA removal was verified by PCR. Ribosomal RNA was depleted using the Ribo-Zero rRNA Removal Kit for bacteria (Illumina). Strand-specific RNA-seq libraries were prepared using the NEBNext Ultra Directional RNA Library Prep Kit for Illumina and sequenced on a GeneMind SURFSeq 5000 sequencing platform.

Raw paired-end reads were quality assessed and trimmed using fastp v0.20.1 (61). Reads were aligned to the PAO1 genome (NC_002516.2) using Bowtie2 v2.4.4 (62) with default settings. Alignments were processed with SAMtools v1.13 (63), including mate correction, sorting, duplicate marking and indexing. Gene-level counts were obtained with featureCounts v2.0.3 (64) in reverse-stranded paired-end mode (-t CDS -s2 -p -B -P -C -Q 10 -D 1000 -M --fraction). Duplicate reads were retained. Counts were supplied to edgeR v4.8.2 (65). Twelve specified rRNA loci were excluded, and lowly expressed genes were filtered before TMM normalization. Each CRISPRi knockdown and the TOB-treated control were compared with the induced non-targeting control. Genes with Benjamini–Hochberg adjusted FDR < 0.05 and |log₂FC| ≥ 1 were considered differentially expressed. The raw sequencing data are deposited in GEO under accession GSE349001 (accessed 1.10.2026, https://www.ncbi.nlm.nih.gov/geo/query/acc.cgi?acc=GSE349001).

### Bioinformatic analyses and data/code availability

Functional enrichment of recurrent gene–aminoglycoside interactions was assessed using KEGG, annotations obtained from the Pseudomonas Genome Database, (66). GO annotations for *P. aeruginosa* PAO1 were consolidated from the Pseudomonas Genome Database, UniProt and QuickGO. Genes annotated to specific GO terms were also assigned to the corresponding broader terms in the GO hierarchy (67). GO analyses used clusterProfiler to identify enriched functions among upregulated and downregulated genes and coordinated expression changes across the full gene list (68). Changes in the activity of coordinately expressed gene groups (iModulons) were analysed using the PRECISE411 reference (69) with the use of PyModulon (70). Guide-efficacy model development and evaluation are outlined in Supplementary Methods. Raw data, processed datasets, annotations, and analysis scripts are available at GitHub: (https://github.com/akawalek-ibb/pao1_crispri_aminoglycoside_analysis), with a final versioned archive to be deposited in Zenodo. Data can also be explored using the CRISPRi browser currently at (https://github.com/Raymi556/CRISPRi-PseudoBrowser), with a website version planned following peer review.

## RESULTS

### Optimized CRISPRi system for gene silencing in *Pseudomonas aeruginosa*

To establish a scalable CRISPRi platform for *P. aeruginosa*, we constructed a dual-vector system in which *dcas9*_Spy_ and sgRNA cassettes were integrated using mini-CTX and mini-Tn7 based vectors, respectively (51, 71) (Fig. 1A). Initial promoter comparisons supported selection of the rhamnose-inducible *rha*p for tightly controlled CRISPRi (Supplementary Fig. S1). In strains carrying the mRFP reporter, rhamnose induction reduced fluorescence 8-to 12-fold with reporter-targeting guides (Fig. 1B) and reporter repression reached a plateau at 0.02% rhamnose (Supplementary Fig. S1A). Targeting selected essential genes impaired growth upon CRISPRi induction (Supplementary Fig. S1B). All three guides targeting the essential DNA translocase gene *ftsK* inhibited growth on solid medium upon CRISPRi induction (Fig. 1C), and in liquid culture, *ftsK*_g1 caused growth inhibition after approximately 2 h, with no corresponding effect in the non-targeting control (Fig. 1D). Next, we next asked whether CRISPRi could reproduce an established antibiotic-response phenotype by targeting *mexXY*, which encodes components of the MexXY-OprM efflux pump, a major determinant of aminoglycoside resistance (72, 73). Repression with four full-match guides sensitized PAO1 to subinhibitory tobramycin (TOB), producing growth phenotypes similar to those of the Δ*mexXY* strain (Fig. 1E). Together, these results validate the CRISPRi system across reporter-repression, essential-gene depletion, and tobramycin-susceptibility phenotypes.

**Figure 1.**
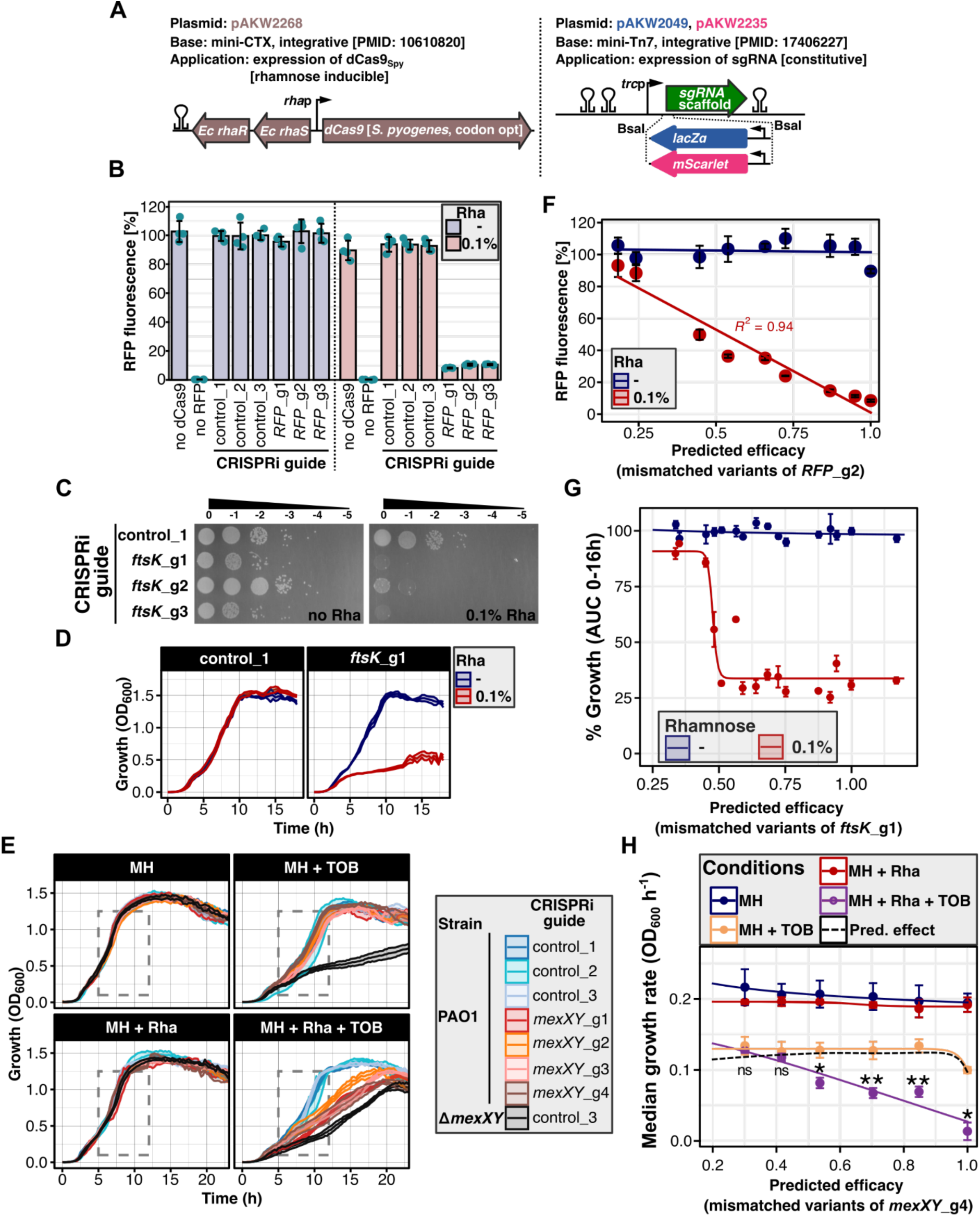
Evaluation of a rhamnose-inducible *S. pyogenes* dCas9-based CRISPRi system in *Pseudomonas aeruginosa* PAO1. **(A)** Architecture of the dual-vector CRISPRi system. A mini-CTX–based vector delivers the rhamnose-inducible *rha*p-*dcas9*_Spy_ cassette, whereas mini-Tn7–based vectors deliver sgRNA-expression cassettes. Spacer insertion by BsaI-mediated Golden Gate assembly replaces a *lacZα* or *mScarlet* cassette in the sgRNA vectors, facilitating identification of spacer-containing clones. **(B)** CRISPRi mediated repression of a reporter gene. PAO1 strains constitutively expressing mRFP and carrying non-targeting or mRFP-targeting guides were grown with or without 0.1% rhamnose (Rha) in LB medium. Data represent mean mRFP fluorescence normalized to OD₆₀₀ ± SD from four biological replicates. Points indicate single measurements. **(C)** Effect of essential-gene repression using CRISPRi. Indicated non-targeting control or *ftsK*-targeting guides were introduced in PAO1 carrying *rha*p_*dcas9*_Spy_ cassette, and growth was assessed on LB agar with gentamicin and triclosan, with or without Rha. **(D)** Effect of *ftsK* repression on bacterial growth in liquid culture. PAO1 strains carrying the non-targeting control_1 or *ftsK*_g1 were grown with or without Rha. Curves represent mean OD₆₀₀ ± SD from four biological replicates. **(E)** Effect of *mexXY* repression on tobramycin susceptibility. PAO1 Δ*mexXY* and CRISPRi strains carrying non-targeting or full-match *mexXY*-targeting guides were grown in MH containing the indicated combinations of 0.1% Rha and subinhibitory 0.4 μg/mL tobramycin (TOB). Curves represent mean OD₆₀₀ ± SD from four biological replicates. Grey dashed lines highlight the region where growth differences are most pronounced. **(F)** Mismatch-based tuning of reporter repression. Single-nucleotide mismatch variants of an mRFP-targeting guide *RFP*_g2, spanning a range of predicted silencing efficiencies, were tested as in (**B**). mRFP fluorescence normalized to OD₆₀₀ is plotted against predicted guide efficacy. Points and error bars represent mean ± SD from four biological replicates. **(G)** Mismatch-based modulation of the growth phenotype associated with *ftsK* repression. PAO1 strains carrying the full-match *ftsK*_g1 guide or its single-nucleotide mismatch variants were grown with or without Rha. Growth was summarized as the area under the OD₆₀₀ growth curve and plotted against predicted guide efficacy. Points and error bars represent mean ± SD from four biological replicates. Mismatch sequences and individual growth curves are provided in Supplementary Fig. S2. **(H)** Tuning tobramycin susceptibility using mismatch variants of the *mexXY*-targeting guide. Growth of strains carrying the full-match *mexXY*_g4 guide or its single-nucleotide mismatch variants was measured in MH medium containing indicated combinations of Rha and TOB. Median growth rates within the selected growth window (Supplementary Fig. S3) were plotted against predicted guide efficacy. Points and error bars represent mean ± SD across four biological replicates. Statistical significance was assessed using a one-sample t-test comparing observed combined-treatment growth rates with the multiplicative expectation calculated from the corresponding single-treatment conditions. * *p* < 0.05, ** *p* < 0.01, ns - not significant.

Because reporter repression plateaued at low rhamnose concentrations (0.02%; Supplementary Fig. S1A), we used spacer mismatches at a fixed inducer concentration to achieve graded knockdown of essential genes without complete growth arrest (8, 41). Mismatched variants of the mRFP-targeting guide produced graded repression strongly associated with predicted guide efficacy (Fig. 1F). Concomitantly, mismatched *ftsK_*g1 variants triggered a threshold-like response, with strong growth inhibition above a predicted efficacy of ∼0.5 (Fig. 1G; Supplementary Fig. S2A-B). By contrast, variants targeting the *lptF–lptG* (PA3828–PA3827) essential operon produced a broader range of intermediate growth defects (Supplementary Fig. S2C–E), suggesting target or guide dependence (8, 29). To determine whether mismatches could also tune antibiotic responses, we tested variants of *mexXY*_g4. The variants produced an efficacy-dependent reduction in growth during TOB exposure (Fig. 1H; Supplementary Fig. S3). Together, these results establish mismatch CRISPRi as an approach for titrating repression and examining its effects on growth and antibiotic susceptibility in *P. aeruginosa*.

To assess applicability beyond PAO1, we targeted essential genes in PA14 and LESB58. CRISPRi induction similarly inhibited growth in both strain backgrounds (Supplementary Fig. S4A). Additionally, to expand the available delivery options, we combined the *dcas9*_Spy_ and sgRNA-expression cassettes on a single mini-Tn7 vector. Upon *ftsK* targeting in PAO1, this construct produced rhamnose-dependent growth inhibition comparable to that of the dual-vector system (Supplementary Fig. S5). Such single-vector configuration may enable CRISPRi in strains restricted to Tn7-mediated integration. Together, these results demonstrate inducible CRISPRi across multiple *P. aeruginosa* backgrounds and provide alternative vector configurations for system delivery.

### Application of CRISPRi-seq to genome-wide fitness profiling in PAO1

In pooled CRISPRi screens, each sgRNA spacer serves as a molecular barcode, allowing individual knockdown abundances to be tracked by sequencing. To determine whether mini-Tn7-mediated delivery could support introduction of complex guide libraries, we quantified vector integration efficacy in PAO1, PA14 and LESB58. Integration frequencies reached approximately 10⁻³–10⁻² integrants per recipient cell in PAO1 and PA14 and approximately 10⁻⁴ in LESB58 (Supplementary Fig. S4B), supporting efficient library delivery across these strains.

To analyse essential and fitness-associated genes in PAO1, we designed an 83,078-sgRNA library targeting 5,557 genes, with dense coverage of 374 genes previously classified as essential, including 320 core-essential genes (14), and 1,035 fitness-associated genes (Fig. 2A; Supplementary Table S2). These groups received up to 28 and 10 full-match guides per gene, respectively, along with mismatched variants, whereas the third category (4,148 genes) was targeted with mostly two full-match guides. In total, the library comprised 26,602 full-match guides, 54,783 mismatch variants and 1,693 non-targeting controls (Supplementary Table S3). Following cloning into pAKW2049 and delivery to PAO1, 82,056 guides (98.8%) were detected in at least one T0 sample, and 79,745 (96.0%) met the cloning criterion of mean abundance ≥10 reads in at least one experimental batch. Only 10 genes in the third category lacked cloned guides (Supplementary Fig. S6; Supplementary Table S4). Thus, the cloned library retained the intended dense coverage of prioritized essential and fitness-associated genes, providing a robust basis for pooled screening.

**Figure 2.**
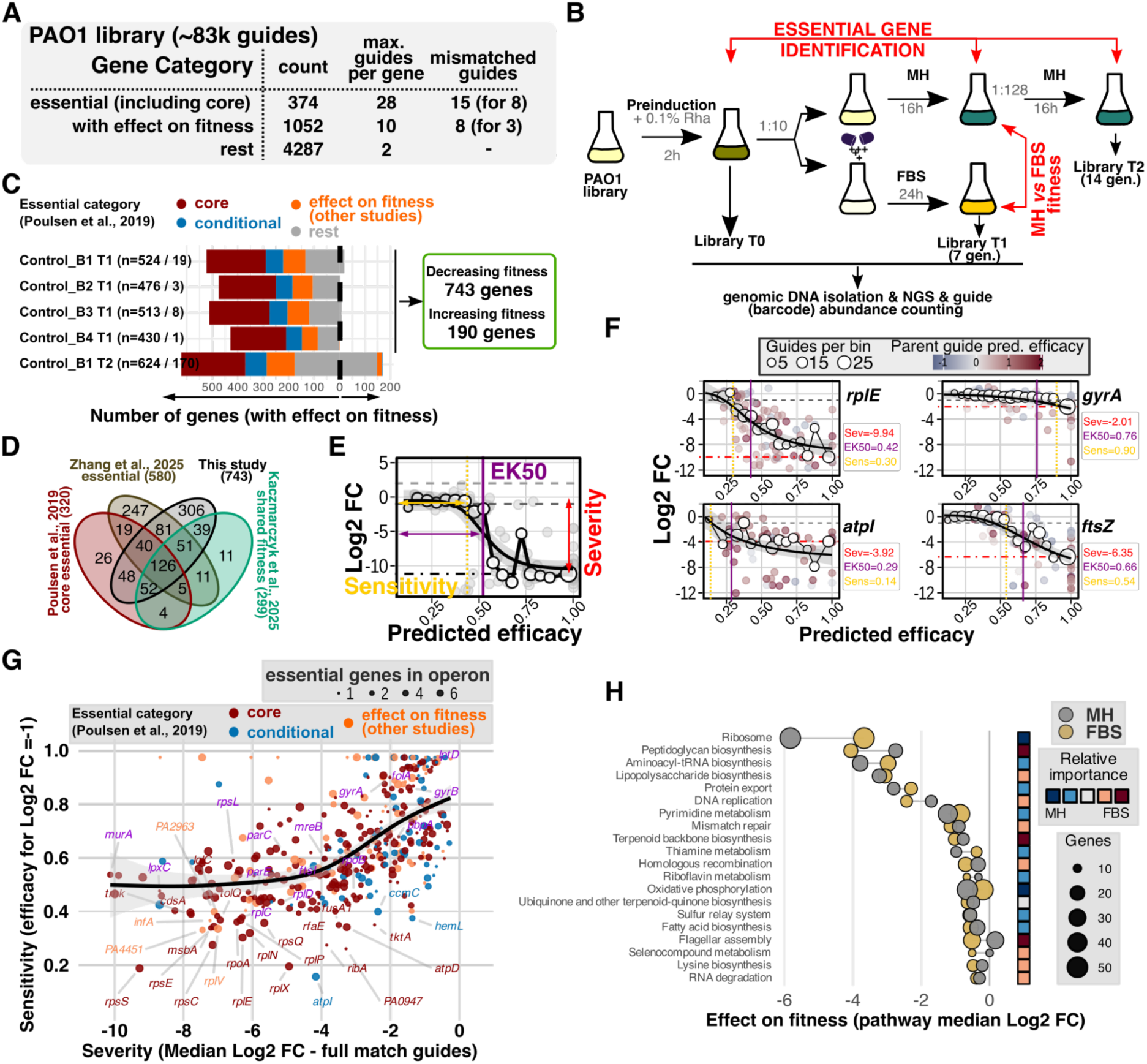
Pooled CRISPRi screening quantifies fitness responses to gene repression in *Pseudomonas aeruginosa* PAO1. **(A)** Schematic of the PAO1 CRISPRi library (∼83k sgRNAs), showing the targeted gene categories, number of full-match guides and mismatch variants. **(B)** Experimental workflow for pooled CRISPRi-seq. Fitness effects were determined by comparing sgRNA abundance after CRISPRi pre-induction (T0) with abundance after approximately 7 (T1) or 14 (T2) additional generations in MH medium. Environmental effects were assessed by comparing induced cultures grown in MH and FBS. **(C)** Numbers of genes showing significant negative fitness effects after CRISPRi induction across independent experimental batches, classified according to previously defined essentiality categories. Numbers on the right indicate the total numbers of unique genes showing significantly decreased or increased fitness in at least one batch or time point. **(D)** Overlap of negative-fitness genes identified here with core-essential genes from Poulsen et al. (14), the PA14 CRISPRi set from Zhang et al. (45), and the PAO1 shared-fitness set from Kaczmarczyk et al. (46). The latter comprises genes with negative-fitness effects across LB, glucose and succinate media. PA14 loci were mapped to PAO1 orthologues. **(E)** Quantitative measures of the fitness response to gene repression. Severity represents the maximal fitness defect associated with repression by full-match guides. Sensitivity represents the predicted guide efficacy at which a sustained fitness effect appears (median log₂FC ≤ −1 for negative effects or ≥ +1 for positive effects). EK_50_ (effective knockdown 50%) represents the predicted guide efficacy producing half of the maximal fitted fitness effect. **(F)** Examples of relationships between predicted sgRNA efficacy and fitness effects for selected genes, illustrating estimation of severity, sensitivity and EK_50_. Data represent the control T1 versus T0 comparison from Batch 1. **(G)** Relationship between CRISPRi severity and sensitivity for genes in libraries grown in MH. Each point represents one gene and is colored according to the essentiality category. Point size indicates the number of genes previously classified as essential by Tn-seq within the corresponding transcriptional unit (Supplementary Table S2). Selected known antimicrobial targets and genes with extreme values are labelled. The black line and shading indicate a generalized additive model fit and 95% confidence interval. Severity and sensitivity represent medians of available estimates across the four untreated T1 versus T0 comparisons (B1–B4). The GAM fit had an effective degrees of freedom (edf) of 3.56 (*p* < 2 × 10⁻¹⁶). **(H)** Pathway-level CRISPRi fitness effects in MH and FBS. Points indicate the median gene-level log₂FC relative to T0 for genes assigned to each KEGG pathway, with point size proportional to the number of genes represented. Displayed pathways were selected according to the largest absolute median fitness effect observed in either condition. Squares summarize gene set enrichment analysis, with color reflecting the normalized enrichment score (NES), indicating whether genes within a pathway are preferentially shifted toward stronger fitness defects in MH or FBS.

To identify repression-associated fitness effects, we propagated the library in MH medium with CRISPRi induction for seven generations in four independent experiments (batches B1–B4) and for 14 generations in B1 (Fig. 2B). Full-match guide abundances at seven (T1) and 14 (T2) generations were compared with T0. Negative-fitness hits required a median adjusted log₂ fold change ≤ −1 across full-match guides and a Stouffer-combined *p* < 0.05 in at least one T1/T0 or T2/T0 comparison. We identified 743 such genes (Fig. 2C; Supplementary Fig. S7; Supplementary Table S5), of which 608 (81.8%) were detected at T1. Concomitantly, guides targeting 190 genes were enriched, predominantly at T2, with many associated with prophage or type IV pilus biogenesis, suggesting a competitive advantage following their repression. Negative-fitness hits overlapped with the core essential sets of Poulsen et al. (14), PAO1 set from Kaczmarczyk et al. (46), and the PA14 CRISPRi screen of Zhang et al. (45) (Fig. 2D). Interestingly, none of the three CRISPRi screens recovered 26 core essential genes (Supplementary Table S5). Overall, pooled CRISPRi captured a broad range of repression-associated fitness effects in PAO1.

### Sequence determinants of CRISPRi guide efficacy

Extensive full-match coverage of fitness-associated genes enabled us to examine determinants of CRISPRi guide efficacy. Because guides targeting the same gene produced different fitness effects, we examined associations with target position, predicted off-targeting and sequence context. Guide depletion weakened with increasing distance from the start codon over the first ∼95 bp, with little further positional association downstream (Supplementary Fig. S8A). In contrast, predicted off-target burden based on seed-sequence similarity (8, 41) was not significantly associated with relative guide depletion (Supplementary Fig. S8B). To examine sequence determinants of guide performance more comprehensively, we trained a lasso model incorporating spacer, variable N position of the PAM, and downstream sequences (Supplementary Fig. S8C). Notably, cytosine (C) at the variable PAM position and pyrimidines (C/T) at the PAM-proximal spacer nucleotide and first downstream base were associated with increased predicted guide efficacy (Supplementary Fig. S8C). Overall the model predictions correlated with observed guide effects for genes excluded from training (median Spearman ρ > 0.5; Supplementary Fig. S8D–F). To compare the model against existing predictors, we scored experimentally tested PAO1 guides using CRISPRi-Pasteur (53) and the mixed-effect random forest (MERF) predictor implemented in the CIAO web server (74). Across 460 genes, the models showed broadly similar ability to rank guides targeting the same gene (Supplementary Fig. S8G,H). To support guide selection in PAO1, PA14 and LESB58, we prioritized candidate sgRNAs by target position and our *P. aeruginosa*-trained model score, with CRISPRi-Pasteur and CIAO predictions provided for comparison (Supplementary Table S6). Concomitantly, predictions for full-match library guides may also help interpret differences in observed guide effects in this study (Supplementary Table S3). Together, these analyses show that CRISPRi guide performance varies with target position and local sequence context.

### Quantifying the fitness effects of essential-gene repression

To move beyond binary essentiality, we analysed gene vulnerability by quantifying fitness defects across a gradient of guide efficacies using data from mismatched sgRNAs. For each gene, we derived three complementary measures (Fig. 2E,F). Severity was the median adjusted log₂ fold change of full-match sgRNAs, reflecting the fitness defect under full-match repression. Sensitivity was the predicted guide efficacy at which the median fitness effect showed a sustained decrease to log₂FC ≤ −1, and EK₅₀ (effective knockdown 50) was the predicted guide efficacy producing half the fitted fitness effect at efficacy 1, assigned for full-match guides (34). Severity and sensitivity were correlated (Spearman ρ = 0.61; Fig. 2G; Supplementary Table S5). Genes with weak full-match fitness defects were rarely highly sensitive to partial repression, whereas genes with strong defects showed a broad sensitivity range. Severity and EK_50_ showed a similar association (Supplementary Fig. S9A). Interestingly, established *P. aeruginosa* antimicrobial targets displayed heterogeneous combinations of severity and sensitivity. Because CRISPRi can repress downstream genes within transcriptional units (TUs) (28, 44), we examined associations with transcriptional-unit organization. Genes in TUs containing more previously classified essential genes (based on Tn-seq screens) showed stronger fitness defects (Supplementary Fig. S9B). The association with sensitivity was weaker, and no such association was detected for EK_50_ (Supplementary Fig. S9C,D). This suggests that operon organization influences fitness-defect magnitude.

We next examined whether the fitness effects of gene repression differed between MH medium and fetal bovine serum (FBS). Stouffer-based and MAGeCK robust rank aggregation analyses identified 269 and 178 genes, respectively, whose repression had significantly different fitness effects between the two media (Supplementary Table S7). Both methods identified stronger defects in FBS following targeting of cell-division genes (*ftsA*, *ftsZ* and *ftsQ*) and *pvdE*, which encodes an ABC transporter required for pyoverdine production. Conversely, targeting purine-biosynthesis genes (*purD*, *purE*, *purL* and *purM*) caused stronger defects in MH. Overall, severity was broadly conserved between media, with similar patterns for sensitivity and EK₅₀ (Supplementary Fig. S10A–C). Because severity could be calculated for more genes than sensitivity or EK₅₀, we used it as our primary vulnerability measure.

To identify biological processes most affected by repression, we compared fitness effects at the KEGG pathway level. The strongest defects involved the ribosome, peptidoglycan biosynthesis, aminoacyl-tRNA biosynthesis, lipopolysaccharide biosynthesis and protein export (Fig. 2H; Supplementary Table S7). Gene set enrichment analysis also identified MH–FBS differences involving the ribosome, flagellar assembly and peptidoglycan biosynthesis. While short-term screening may not capture all conditionally essential functions, these results show that CRISPRi effectively reveals environment-dependent gene and pathway vulnerabilities.

### CRISPRi-seq-based identification of genes modulating aminoglycoside susceptibility

To probe how CRISPRi-mediated repression affects antibiotic susceptibility, we exposed the pooled library to four concentrations of tobramycin (TOB; MIC = 1 µg mL⁻¹) spanning 0.025–1× MIC and amikacin (MIC = 4 µg mL⁻¹) at 0.5× MIC (Fig. 3A). Sensitizing and protective interactions were defined by significant depletion and enrichment, respectively, relative to induced, untreated controls, using condition-specific thresholds (Supplementary Table S8). Interaction counts and directionality varied across treatments, with predominantly protective calls at 1× MIC TOB (Fig. 3B; Supplementary Fig. S11). Comparison with the initial library (T0) showed that knockdowns strongly depleted in untreated controls were relatively retained at 1× MIC TOB, suggesting that differential growth could contribute to apparent protective interactions (Supplementary Fig. S12A). Longer TOB exposure did not increase the number of interactions (T2; Supplementary Table S8). Mismatch-guide analysis identified eight additional sensitizing candidates with effects at partial predicted guide efficacies that were not captured by full-match gene-level calls (Supplementary Fig. S13; Supplementary Table S8).

**Figure 3.**
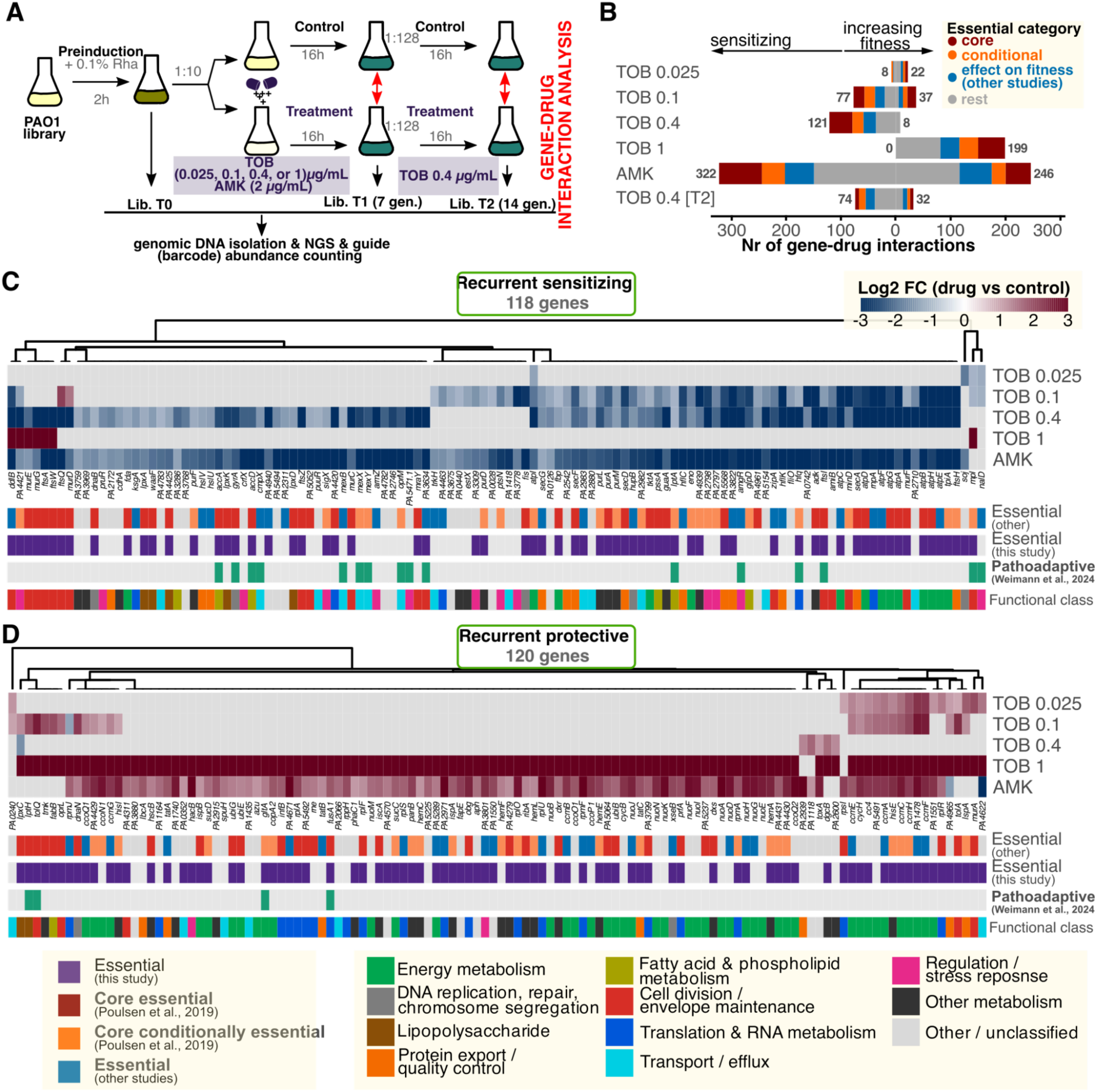
High-throughput CRISPRi-seq identifies genetic determinants of aminoglycoside susceptibility. **(A)** Schematic overview of the pooled CRISPRi-seq experiment and the comparison used to identify gene–drug interactions. Following CRISPRi induction, sgRNA abundance under aminoglycoside treatment was compared with the corresponding untreated induced libraries. **(B)** Numbers of significant sensitizing and protective gene–drug interactions identified in the indicated comparisons. Interactions were defined using a Stouffer-combined *p* < 0.01 together with condition-specific median log₂ fold-change thresholds (Supplementary Fig. S11). Bars are subdivided according to the essentiality category. **(C–D)** Interaction-severity heatmaps for recurrent sensitizing **(C)** and recurrent protective **(D)** gene–aminoglycoside interactions. Recurrence was defined as a significant interaction in the same direction in at least two of the five independent T1 aminoglycoside comparisons. Cell values represent the median adjusted log₂ fold change for the corresponding treatment-versus-control comparison. Nonsignificant interactions are masked with grey color. Genes were clustered by their interaction patterns across conditions. Annotation rows indicate published essentiality, fitness defects identified in this study, pathoadaptive status according to Weimann et al. (75), and manually reviewed functional classes based on KEGG and product annotations.

Overall, across the five T1 conditions, the full-match based analysis identified 118 recurrent sensitizing and 120 recurrent protective genes, defined by same-direction interactions in at least two conditions (Fig. 3CD; Supplementary Table S8). Analysis of the same data using MAGeCK robust rank aggregation supported 68 sensitizing and 69 protective genes in the same direction in at least two T1 conditions (Supplementary Fig. S12B; Supplementary Table S8). Comparison with a previous aminoglycoside Tn-seq screen by Murray et al., (24) matched 28 recurrent sensitizing and 11 recurrent protective hits (Supplementary Table S8). Subsequent analyses focused on sensitizing interactions to identify candidate targets that may enhance antibiotic activity.

The recurrent sensitizing set was enriched in genes whose repression also impaired fitness without antibiotic exposure (79/118; Fisher’s exact test, *P* = 7.3 × 10⁻⁴²). Strikingly, for a subset of sensitizing genes, fitness defects emerged at lower predicted guide efficacies during TOB or AMK exposure, consistent with increased vulnerability to partial repression (Supplementary Fig. S14). The 118 sensitizing genes mapped to 79 TUs (Supplementary Table S8), and represented various biological processes. Lipopolysaccharide biosynthesis, protein export and purine metabolism remained enriched when each transcriptional unit was counted once (Supplementary Table S8). Peptidoglycan biosynthesis and oxidative phosphorylation were enriched only when genes were counted individually, with hits concentrated in two TUs and the ATP synthase TU, respectively. The sensitizing set also included 16 previously reported pathoadaptive genes ((75); Fig. 3C), including *hfq, accD, ftsB* (PA3634), *gyrA, accA, lptA, mpl* and *ftsI*. Repression of these genes also impaired fitness in untreated medium, possibly linking functions implicated in adaptation during infection with growth and aminoglycoside responses.

### Partial repression of selected essential genes enhances tobramycin activity

To validate screen-derived interactions, we selected *atpI* (ATP synthase protein I), *secA* (Sec translocation ATPase), and *ftsZ* (cell-division GTPase), all of which exhibited efficacy-dependent sensitization in our screen (Supplementary Fig. S15A-C). We also selected *accC*, encoding the biotin carboxylase subunit of acetyl-CoA carboxylase, based on sensitization observed with mismatch guides targeting *accC* and *accB* and the identification of *accA* and *accD* as recurrent sensitizing genes (Supplementary Figs. S13 and S15). For each target, we tested one full-match guide and five mismatch variants spanning a range of predicted efficacies. Median growth rates under combined CRISPRi induction and TOB treatment were compared with the multiplicative expectation from the individual treatments (Fig. 4A; Supplementary Fig. S16). Ultimately, sensitizing interactions were confirmed for all four targets (Fig. 4B–E). For *atpI*, TOB-dependent sensitization was observed across all six guides. In contrast, for *accC*, *secA* and *ftsZ*, sensitization was detected only with two mismatched variants of intermediate predicted efficacy per gene, showing that partial repression can reveal enhanced TOB-mediated growth inhibition in individual assays.

**Figure 4.**
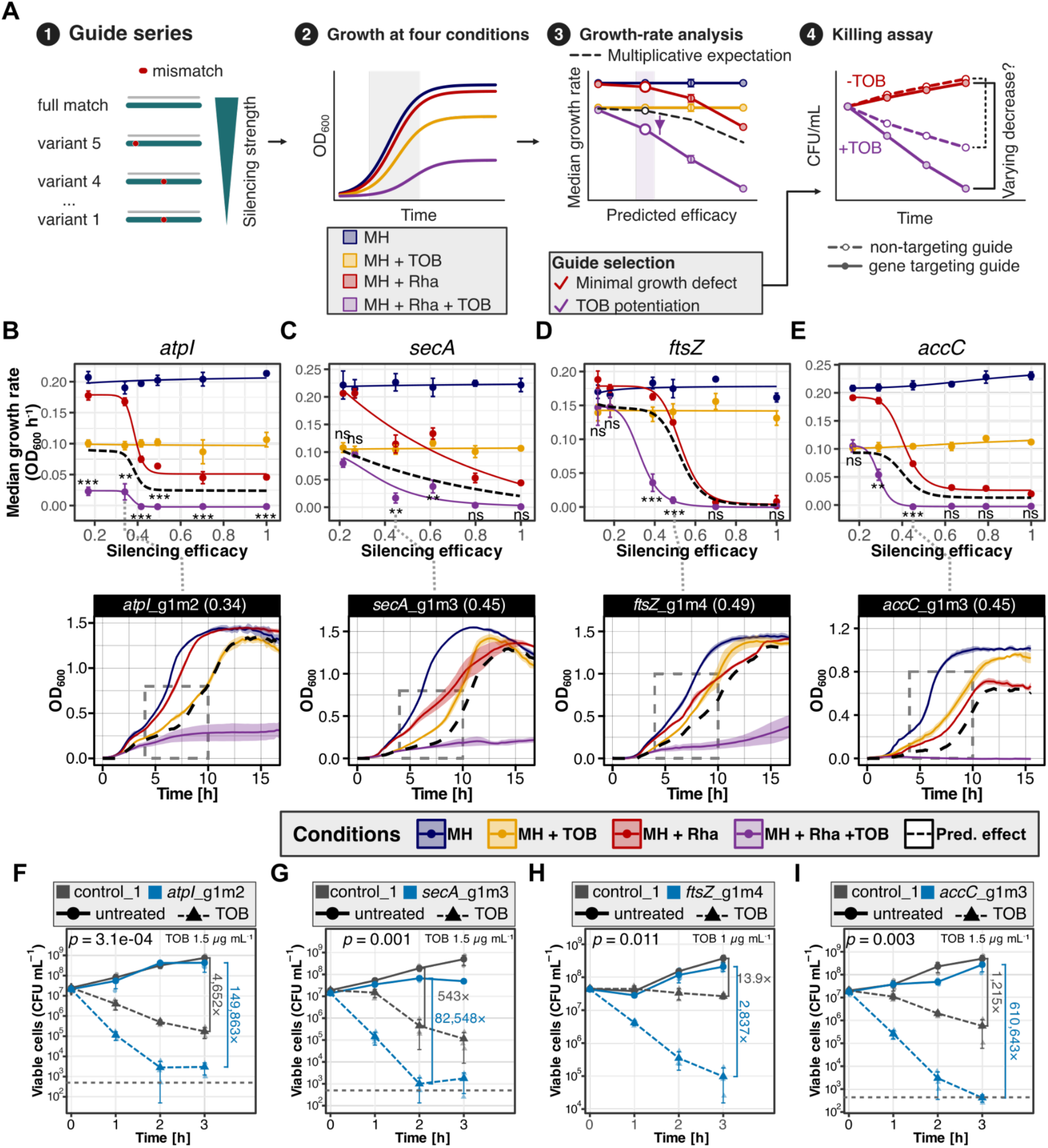
Experimental validation of selected essential gene–aminoglycoside interactions. **(A)** Schematic of the validation workflow. Strains harbouring CRISPRi guide series spanning a range of predicted silencing efficiencies were tested in four growth conditions. Median growth rates were used to test whether the combined effects of gene repression and TOB differed from those expected from their individual effects (multiplicative model). One guide per gene was then selected to assess tobramycin-mediated killing. **(B–E)** Growth-based validation of the effects of *atpI* (**B**), *secA* (**C**), *ftsZ* (**D**) and *accC* (**E**) repression on TOB activity. For each gene, strains carrying a full-match guide or mismatch variants were grown in MH medium, MH containing 0.4 μg/mL TOB, MH containing 0.1% rhamnose (Rha), or MH containing both Rha and TOB. Median growth rates within the selected growth window (dashed lines, see Supplementary Fig. S16) were plotted against predicted guide efficacy. Points and error bars represent mean ± SD across four biological replicates. The black line indicates the multiplicative expectation calculated from the corresponding single-treatment conditions. A one-sample, one-sided *t*-test assessed whether observed combined-treatment growth rates were lower than the multiplicative expectation. Sensitizing interactions required *p* < 0.05 and an observed growth rate below the multiplicative expectation by at least 10% of the untreated growth rate. Significance: * *p* < 0.05, ** *p* < 0.01, *** *p* < 0.001; ns, not significant. For each gene, growth curves for the selected guide are shown at the bottom. Growth data indicate mean OD₆₀₀, shaded areas indicate ± SD, and the black dashed line shows the multiplicative expectation. Numbers in brackets indicate the predicted efficacy of the displayed mismatch guide. **(F–I)** Tobramycin killing assays for PAO1 carrying *atpI*_g1m2 (**F**), *secA*_g1m3 (**G**), *ftsZ*_g1m4 (**H**) or *accC*_g1m3 (**I**), compared with the non-targeting control_1 guide. Strains were precultured in MH with 0.1% rhamnose, and subsequently transferred to the same medium with or without TOB at the concentration indicated in each panel. Viability was determined as CFU mL⁻¹ at the indicated time points. Data represent mean ± SD from three biological replicates, with shapes indicating individual replicate values. Selected timepoint fold decreases indicate the reduction in viable counts associated with TOB treatment. The dashed line indicates the detection limit of 500 CFU/ml. Statistical significance was assessed by comparing biological-replicate matched log₁₀ CFU reductions between the control and knockdown strains using a paired, two-sided *t*-test, and *p*-value is reported.

To determine whether growth restriction in presence of TOB translated into enhanced killing, we tested PAO1 strains carrying *atpI*_g1m2, *secA*_g1m3, *ftsZ*_g1m4 or *accC*_g1m3, which retained measurable growth under CRISPRi induction alone. Following inducing preculture, knockdown strains and the non-targeting control were exposed to TOB and viability was measured by CFU enumeration. All four knockdowns significantly enhanced TOB killing (Fig. 4F–I). At the respective assay endpoints, survival fractions were approximately 32-, 152-, 204- and 503-fold lower than those of matched non-targeting controls for *atpI*, *secA*, *ftsZ* and *accC*, respectively. The corresponding *atpI*, *secA* and *ftsZ* guides also sensitized PA14 to TOB (Supplementary Fig. S17). Together, these results show that partial repression of distinct cellular functions can enhance both TOB-mediated growth inhibition and killing.

### Transcriptional and permeability assays reveal distinct consequences of selected knockdowns

To verify target repression and examine its transcriptional consequences, we performed RNA-seq on induced CRISPRi strains carrying the mismatch guides selected for killing assays (Supplementary Table S9). Relative to the induced non-targeting control, *atpI, accC, secA* and *ftsZ* transcript levels were reduced to approximately 25%, 52%, 42% and 42%, respectively (Fig. 5A–D). Targeting *atpI* reduced expression throughout the *atpIBEFHAGDC* operon. By contrast, *accC* repression was comparatively localized, without co-repression of the upstream *aroQ1* and *accB* genes. Targeting *ftsZ* also reduced expression of downstream *lpxC* and *PA4406.1*. Targeting *secA* was accompanied by reduced expression of neighbouring genes not predicted to belong to the same operon. Thus, all four constructs reduced target expression, with differing local transcriptional effects.

**Figure 5.**
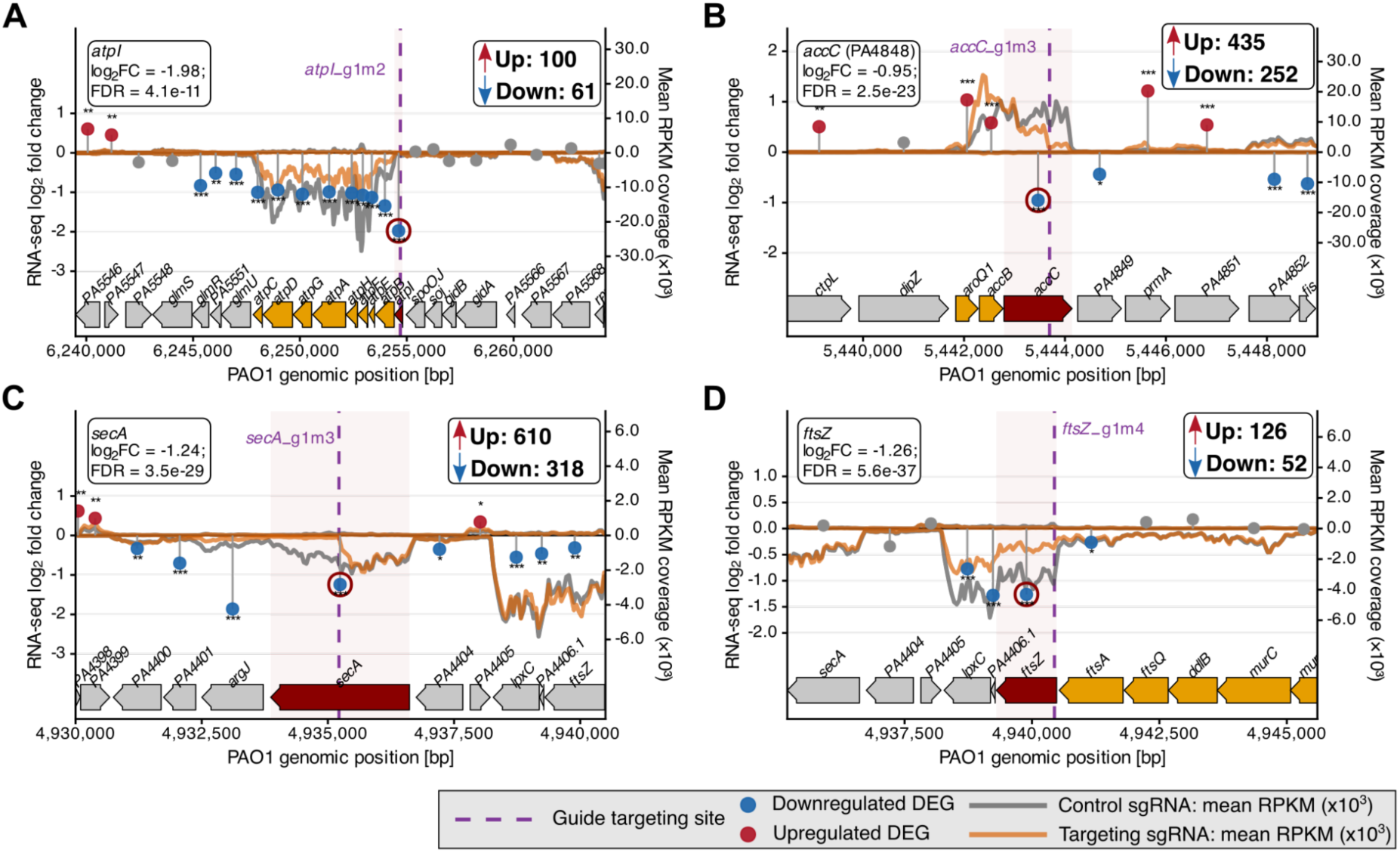
Local transcriptional effects of CRISPRi-mediated gene repression of *atpI*, *accC*, *secA* and *ftsZ*. **(A–D)** RNA-seq coverage and differential expression around regions targeted by *atpI*_g1m2 (**A**), *accC*_g1m3 (**B**), *secA*_g1m3 (**C**) and *ftsZ*_g1m4 (**D**). Each knockdown was compared with the induced strain carrying the control_1 guide. Coverage shows mean RPKM ± SD from three biological replicates, with positive and negative values representing forward- and reverse-strand reads, respectively. Target genes are shown in red, other genes in the same annotated transcriptional unit in orange, and neighbouring genes in grey. Violet dashed lines mark guide target sites. Points show gene-level log₂ fold changes relative to the control strain. Asterisks indicate FDR-adjusted *p* values: * < 0.05, ** < 0.01 and *** < 0.001. Up and Down indicate genome-wide counts of upregulated and downregulated genes, respectively, at FDR <0.05 and |log₂FC| ≥1.

Global responses differed markedly among knockdowns. At FDR < 0.05 and |log₂FC| > 1, *accC, secA, ftsZ* and *atpI* repression altered 687, 928, 178 and 161 genes, respectively (Fig. 5; Supplementary Table S9). TOB exposure alone altered expression of 64 genes, 53 of which were among the genes affected by at least one knockdown (Supplementary Table S9). We next compared transcriptional responses among recurrent sensitizing and protective genes (Fig. 6A,B). Using FDR <0.05 without a fold-change cutoff, 23 sensitizing and 21 protective genes showed expression changes in at least three knockdowns, including three and four genes, respectively, in all four. Interestingly, *accC* and *secA* repression produced several shared transcriptional signatures. Similarly, transcriptome projection onto PRECISE411 iModulons indicated higher RpoS-2 and quorum-sensing-associated activity and lower Translational-1 activity in the *accC* and *secA* knockdowns (Fig. 6C). Consistently, gene ontology (GO) analysis identified enrichment of translation-related genes among downregulated genes in both knockdowns (Fig. 6D). ATP synthase expression was also reduced: all eight structural genes (*atpBEFHAGDC*) were significantly downregulated after *accC* repression, and all nine *atpIBEFHAGDC* genes after *secA* or direct *atpI* targeting (Fig. 6A), whereas ATP synthase repression was not observed after *ftsZ* targeting. Other responses were target dependent. *secA* repression increased PvdS, PchR and iron-acquisition signatures, accompanied by upregulation of iron-import and siderophore-biosynthesis genes. In contrast, *accC* repression downregulated iron-import genes and increased HexR-1 activity (Fig. 6C). Fatty-acid-biosynthesis genes were enriched among upregulated genes after *accC* repression, suggesting a compensatory response. Genes encoding outer-membrane proteins were overrepresented among genes downregulated after *accC* repression and upregulated after *secA* repression. To examine whether these envelope-associated responses accompanied altered outer-membrane permeability, we measured NPN fluorescence. Increased fluorescence was observed in the tested *accC* and *secA* knockdowns, whereas no comparable increase was detected for *atpI* or *ftsZ* under the tested conditions (Fig. 6E). These results are consistent with altered outer-membrane permeability in a subset of sensitized strains. Together, these findings suggest that sensitization may be multifactorial, with distinct transcriptional responses across knockdowns and altered outer-membrane permeability in the *accC* and *secA* strains.

**Figure 6.**
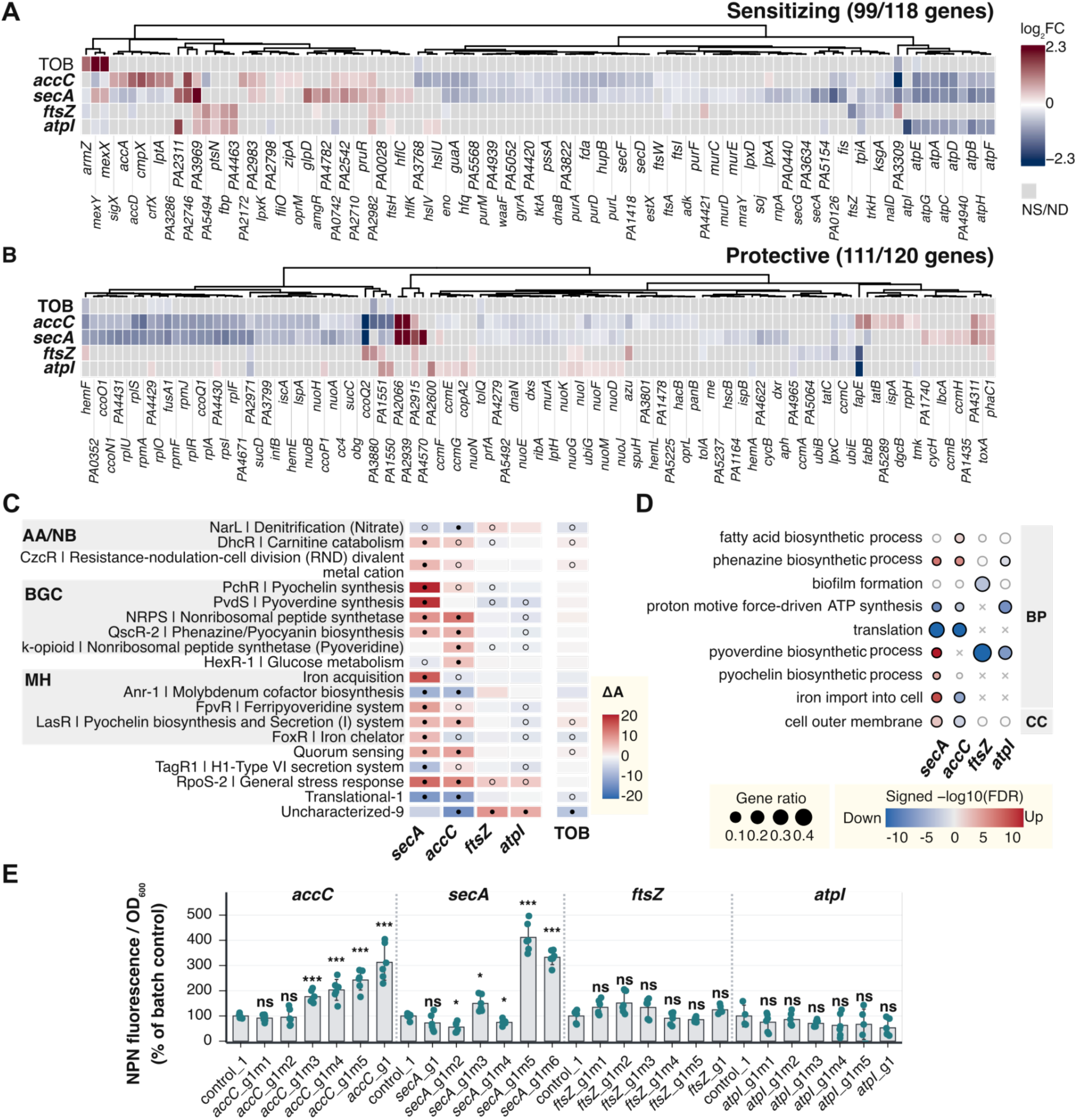
Global transcriptional responses and outer-membrane permeability following CRISPRi-mediated gene repression of selected essential genes. **(A–B)** RNA-seq expression changes in genes belonging to the recurrent sensitizing (**A**) and protective (**B**) sets. The TOB rows indicate the response of the non-targeting control strain to TOB exposure. Colours indicate log₂ fold changes relative to the corresponding induced non-targeting control without TOB. Changes with FDR < 0.05 are coloured. Nonsignificant cells are masked in grey. **(C)** Changes in iModulon activity following the indicated knockdowns or TOB treatment, relative to the induced non-targeting control. An iModulon represents a set of genes with coordinated expression identified by independent component analysis of transcriptomic data. iModulon activity changes summarize the coordinated transcriptional responses within each module (69). iModulons with |Δactivity| >5 in at least one comparison are shown, grouped using PRECISE411 annotations (69). Data for all are presented in Supplementary Table S9. AA/NB, amino acid and nucleotide biosynthesis; BGC, biosynthetic gene cluster; MH, metal homeostasis. **(D)** Selected GO terms enriched among upregulated or downregulated genes. Bubble size indicates the fraction of differentially expressed genes included in each GO analysis that are associated with the indicated term. Color shows signed −log₁₀(FDR), with positive values indicating enrichment among upregulated genes and negative values indicating enrichment among downregulated genes. Grey indicates FDR ≥ 0.05. Crosses indicate GO terms not returned by the enrichment analysis (e.g. no DEG). BP, biological process; CC, cellular component. **(E)** Outer-membrane permeability assessed by NPN fluorescence following induction of the indicated CRISPRi strains. NPN fluorescence was divided by OD₆₀₀ and expressed relative to the mean of the corresponding batch control set as 100%. Bars show mean ± SD. Points represent independent biological replicates. Each strain was compared with its batch control using a two-sided Welch’s t-test on log₂-transformed NPN/OD₆₀₀ values, with Holm correction for multiple comparisons. Adjusted *p* values: * <0.05, ** <0.01, *** <0.001; ns, not significant.

## DISCUSSION

Here we developed a rhamnose-regulated CRISPRi platform for *P. aeruginosa* combining color-based guide cloning, sequence-informed guide selection and efficient chromosomal integration of complex libraries. Its compact sgRNA vector facilitates cloning and handling (Fig. 1; Supplementary Fig. S4B). The 83,078-guide library constructed here enabled genome-wide analysis of gene contributions to fitness and aminoglycoside susceptibility in PAO1. This platform complements inducible PAO1 systems (46, 47), doxycycline-inducible CRISPRi-seq in PA14 (45), and a rhamnose-controlled dCas9_Spa_ system with similar dual vector architecture (48). Importantly, rhamnose induction benefits chemical–genetic experiments by avoiding an additional antibacterial compound. Our data also indicate that guide performance was associated with target position and nucleotide context within and around the PAM. The sequence-context associations agree with previous bacterial guide-prediction studies (53, 74). These findings informed guide recommendations for PAO1, PA14 and LESB58 to support individual CRISPRi experiments and compact library design (Supplementary Table S6).

Repression of 743 genes reduced competitive fitness in MH medium, encompassing essential genes and additional growth determinants. Some fitness defects may reflect co-repression of downstream genes within the same transcriptional unit. Translation, aminoacyl-tRNA biosynthesis, envelope synthesis and protein export were among the functions most affected by repression. The relatively high vulnerability of translation and cell-wall functions in PA14 (45), tRNA synthetases in *Mycobacterium tuberculosis* (9) and peptidoglycan/division functions in *Acinetobacter baumannii*(34) suggests shared vulnerabilities across these pathogens. Interestingly, genes within the core-essential set of *P. aeruginosa* genes (14) differed substantially in severity and sensitivity (Supplementary Table S5). This agrees with depletion studies showing that essential targets tolerate different reductions in abundance (7, 76). Full-match guides also differed in performance. Weak fitness defects may therefore reflect tolerance to reduced gene expression or insufficient repression, despite the denser guide coverage of core-essential genes. Within the limited conditions tested, fitness effects were broadly conserved between MH medium and fetal bovine serum (FBS), although a subset of genes showed medium-specific responses to silencing. Overall, these data extend *P. aeruginosa* essential-gene catalogs by quantifying the fitness consequences of gene repression.

The 118 recurrent sensitizing loci included established determinants of aminoglycoside susceptibility such as *ftsH*, *amgR* and *tpiA* (77–79). Several hits, including *lptA*, *ksgA* and *atpI*, also showed concordant aminoglycoside phenotypes in the Tn-seq study of Murray et al. (24), and *lptA* has independently been linked to intrinsic resistance to multiple aminoglycosides (80). The sensitizing effect of *gyrA* repression is also notable in the context of ciprofloxacin–tobramycin synergy in PAO1 and hypermutable strains, although the relationship between these genetic and pharmacological interactions remains unresolved (81, 82). Protective interactions involving *nuo* and *ccm* genes agree with reduced tobramycin susceptibility reported for respiratory-chain and cytochrome maturation mutants (83). These findings fit the established contribution of respiratory state to aminoglycoside uptake and lethality (84). Overall, recovery of known determinants supports the screen, which also extends aminoglycoside-response profiling to essential genes and other growth determinants.

Individual growth and killing assays confirmed enhanced tobramycin activity following partial repression of *atpI*, *accC*, *secA* and *ftsZ* (Fig. 4). RNA-seq confirmed target repression and showed that some guides also reduced expression of neighbouring genes (Fig. 5). Targeting *atpI* repressed the entire *atpIBEFHAGDC* operon, so the phenotype cannot be attributed to *atpI* alone. Guides targeting each of the nine operon genes also produced recurrent sensitizing interactions in the pooled screen, supporting ATP-synthase involvement (Fig. 3C). In PA14, mutations affecting ATP synthase or its promoter have been associated with increased aminoglycoside susceptibility and inner-membrane hyperpolarization (85). Disruption of *atpA* also increases gentamicin susceptibility in *Staphylococcus aureus* (86), while the ATP-synthase inhibitor venturicidin A potentiates gentamicin against *P. aeruginosa* (87). Together, these findings support ATP synthase as a candidate target for aminoglycoside potentiation.

Sensitization following *accC* targeting implicates the acetyl-CoA carboxylase (ACC) complex in aminoglycoside susceptibility, consistent with sensitizing trends across the other genes encoding its subunits. Interestingly, all four ACC subunit-encoding genes were also identified in the pathoadaptive set, although adaptive mutations need not reproduce the effects of gene repression (75). Previous studies demonstrate distinct effects of altering fatty-acid biosynthesis. Treatment with 2-aminooxazole, used as an ACC inhibitor, or triclosan, which inhibits the fatty-acid synthesis enzyme FabI, enhanced ciprofloxacin-mediated killing of resistant *P. aeruginosa* (88). By contrast, glutamate supplementation increased ACC activity and unsaturated-fatty-acid synthesis, increasing membrane permeability, gentamicin uptake and killing, and treatment with either inhibitor reduced this potentiation (89). These findings suggest that the effects of altering fatty-acid biosynthesis depend on the antibiotic, the perturbation and the metabolic context. Increased NPN fluorescence after *accC* repression suggests altered outer-membrane permeability (Fig. 6E), which may contribute to sensitization.

The *secA* phenotype suggests a contribution of Sec-dependent protein export to aminoglycoside susceptibility. In PA14, disruption of *secD*, *yajC* and the translocation-associated chaperone gene *ppiD* reduced tobramycin MICs (90). Because SecA drives protein translocation across the inner membrane, reduced export could impair envelope assembly or protein quality control. However, the substrates and processes responsible for sensitization remain unidentified. Comparison of the *accC* and *secA* knockdowns revealed shared responses alongside important differences (Fig. 6). Both showed increased NPN fluorescence, consistent with increased outer-membrane permeability. Their iModulon projections showed higher RpoS-2 and quorum-sensing-associated activities and lower Translational-1 activity. The reduced Translational-1 activity agreed with enrichment of translation-related genes among downregulated transcripts. These signatures could however reflect growth inhibition or compensatory responses. The iron responses differed: *secA* repression increased PvdS/PchR-associated signatures and iron-acquisition gene expression, whereas *accC* repression downregulated iron-import genes. Both knockdowns also reduced ATP-synthase expression, providing a possible connection to the ATP-operon phenotype discussed above. Together, these findings suggest that sensitization after *accC* and *secA* repression may involve several physiological changes, with altered outer-membrane permeability as one possible contributor.

Sensitization following *ftsZ* targeting occurred without a comparable increase in NPN fluorescence or coordinated repression of ATP-synthase genes, indicating that these responses were not shared by all sensitized strains. Co-repression of downstream *lpxC* and *PA4406.1* prevents attributing the *ftsZ*-targeting phenotype exclusively to FtsZ depletion. However recurrent sensitizing loci included *murF*, *ftsI*, *ddlB* and *mraY* within the same annotated cell-division transcriptional unit. Division-site regulation was also implicated by recurrent sensitizing interactions involving *minD* and sensitization to AMK following *minE* targeting. Previous studies linked *ftsZ* depletion and Min-system deficiency to cell elongation (43, 49), and cell-division defects to increased tobramycin susceptibility (91). Together, these findings implicate cell-division and envelope functions in aminoglycoside susceptibility.

Several aspects of the experimental design and analysis limit interpretation. Denser guide coverage of essential and fitness-associated genes may have increased detection power for these categories. CRISPRi polarity also limits single-gene attribution within transcriptional units (28, 29). Interaction calls also varied between analysis methods. At strongly growth-inhibitory TOB concentrations, fewer population doublings may reduce depletion of guides that impair fitness, producing apparent protective interactions in comparisons with untreated cultures (Supplementary Fig. S12A). Sensitivity and EK₅₀ were estimable for only a subset of genes and relied on predicted guide efficacy rather than directly measured repression. Estimates may also be influenced by parent-guide activity, sequencing depth and the response range captured by each mismatch series. Growth-rate differences may also contribute to the observed transcriptional responses. Membrane potential and intracellular aminoglycoside accumulation were not measured, leaving their contributions to sensitization unresolved. Finally, screening primarily in planktonic PAO1 limits extrapolation to other strain backgrounds and growth conditions.

Despite these limitations, the platform, screening data and accompanying CRISPRi browser provide a resource for prioritizing and testing essential-gene–antibiotic interactions. Browser code is currently available on GitHub, with a website version planned.

## AUTHORS CONTRIBUTIONS

**AK** conceived the study, designed the experimental framework and pooled sgRNA library, performed the pooled CRISPRi-seq and RNA-seq screens, conducted the growth and killing assays, analyzed the data, curated and deposited the data and code, prepared the figures, and wrote the original draft. **WCZ** conducted vector construction, preliminary experiments, growth and killing assays, and NPN permeability measurements. **BD** cloned the pooled sgRNA library and, together with **LFPM** and **FP**, contributed to vector construction and preliminary tests. **FP** also performed the reporter fluorescence assays. **JG** and **KZ** designed the CRISPRi-seq sequencing strategy and performed the sequencing. **REGC** developed the interactive CRISPRi browser. **AAB** and **XL** contributed to data analysis and biological interpretation. **AK**, **WCZ**, **AAB**, and **XL** revised and edited the manuscript. All authors read and approved the final version.

## ACKNOWLEDGEMENTS

This work was funded by the National Science Centre in Poland grant 2021/43/D/NZ2/02151 awarded to A.K.

## SUPPLEMENTARY DATASET

**Supplementary Table S1. Bacterial strains, plasmids, oligonucleotides used in this study**

**Supplementary Table S2. Essentiality and antimicrobial-fitness evidence used to design the PAO1 CRISPRi library.**

**Supplementary Table S3. Sequences, target assignments and activity predictions for the PAO1 guide library.**

**Supplementary Table S4. CRISPRi library representation.**

**Supplementary Table S5. Genetic determinants of competitive fitness in antibiotic-free medium.**

**Supplementary Table S6. Recommended CRISPRi guides for Pseudomonas aeruginosa PAO1, PA14 and LESB58.**

**Supplementary Table S7. CRISPRi gene and pathway effects in MH and fetal bovine serum.**

**Supplementary Table S8. CRISPRi-seq analysis of gene–aminoglycoside interactions.**

**Supplementary Table S9. RNA-seq expression changes, GO enrichment and iModulon analyses following CRISPRi-mediated gene repression or tobramycin exposure.**

## SUPPLEMENTARY METHODS

### CRISPRi library cloning

The pooled oligonucleotide library was synthesized by GenScript as 78-nt oligonucleotides with the structure 5′-GTCTGGAGCGCGGATCTAGGTCTCCTAGT-N20-GTTTAGAGACCCAGGACGCGGTAAAC ACC-3′, where N20 represents the spacer. The pool was amplified with primers #183 and #184 using KAPA HiFi polymerase in parallel PCR reactions, and the 78-bp product was gel-purified using the GeneJET Gel Extraction Kit (Thermo Scientific). Purified products were inserted into pAKW2049 by scaled Golden Gate assembly. Following bulk electroporation into *E. coli* DH5α, approx. 30M transformants were pooled and plasmid DNA was isolated using the Plasmid Midi AX kit (A&A Biotechnology). The plasmid library was subsequently electroporated into *E. coli* S17-1 λpir and transferred to the dCas9-expressing PAO1 recipient by mini-Tn7-mediated chromosomal integration. Approximately 2.5 × 10⁸ PAO1 integrants were recovered, corresponding to approximately 3,000-fold representation of the designed library. Cells were scraped from the plates, pooled and supplemented with glycerol to a final concentration of 16%, then aliquoted, flash-frozen in liquid nitrogen and stored at −80°C.

### CRISPRi-seq library preparation

Genomic DNA was isolated from bacterial pellets using the Genomic Mini kit (A&A Biotechnology, Poland). DNA integrity was assessed by 1% agarose gel electrophoresis, quantified using Qubit Broad Range chemistry on a Qubit 3.0 fluorimeter (Thermo Fisher Scientific), and diluted to 10 ng µL⁻¹. The sgRNA-containing region was amplified using primers carrying Nextera XT adapter overhangs and variable-length heterogeneity spacers to increase sequencing diversity (primers #185-198, Supplementary Table S1). Primer variants were combined at equimolar concentrations and used as 10 µM pools. Each sample was amplified in three technical PCR reactions using 2× Phanta Vazyme PCR Mix (Vazyme) in a final volume of 10 µL containing approximately 10 ng genomic DNA, 5 µL 2× PCR mix, 3 µL MQ water, and 0.5 µL each of the XT_CRISPRiTn7_F and XT_CRISPRiTn7_R primer pools. PCR amplification consisted of 95°C for 3 min, followed by 25 cycles of 98°C for 15 s, 60°C for 10 s and 72°C for 15 s, with a final extension at 72°C for 1 min. Amplicons were verified by agarose gel electrophoresis, technical PCR replicates were pooled, and purified using Mag-Bind TotalPure NGS beads (Omega Bio-tek). Purified amplicons were indexed by a second PCR with Nextera XT index primers (Illumina), purified again with magnetic beads, quantified using the NEBNext Library Quant Kit (New England Biolabs), pooled equimolarly and sequenced on an Illumina NovaSeq 6000 or GeneMind platform including 5% PhiX, targeting approximately 50 million reads per sample.

### CRISPRi-seq data analysis

Guide counts were analysed separately by batch after filtering for mean untreated-T0 abundance ≥100 and applying filterByExpr, followed by TMM normalization across retained guides, robust dispersion estimation and quasi-likelihood testing in edgeR. Guide log₂ fold changes were adjusted by subtracting the median value for non-targeting guides, and gene-level effects (median Log2 FC) were calculated as the median adjusted value of full-match guides. Full-match guide *p* values were combined per gene using Stouffer’s method, and screening calls used the unadjusted combined *p* values. Negative-fitness calls required Stouffer *p* < 0.05 and medLFC ≤ −1, whereas drug interactions required *p* < 0.01 and comparison-specific effect-size thresholds defined by median ± 5 median absolute deviation, with a minimum absolute cutoff of 1 (Supplementary Table S8). Recurrent interactions were significant in the same direction in at least two of the five T1 comparisons between aminoglycoside-treated and matched untreated cultures. The recurrent hit sets were largely preserved when Benjamini–Hochberg-adjusted *p* < 0.05 was additionally required alongside the original screening criteria, retaining 113/118 (95.8%) sensitizing and 111/120 (92.5%) protective hits (Supplementary Table S8). Additionally we used MAGeCK robust rank aggregation (RRA) as a complementary analysis to assess whether gene-level findings depended on the analysis method. MAGeCK combines guide rankings to identify genes supported by consistent guide depletion or enrichment, with gene-level false-discovery-rate correction (59). Full-match guides were also grouped by annotated transcriptional unit to calculate TU-level effects and interaction calls, using median adjusted log₂ fold changes, Stouffer-combined *p* values and effect thresholds calculated from TU-level distributions (Supplementary Table S8).

To quantify gene-level fitness responses across predicted guide efficacies, full-match and mismatch guides from all parent-guide families targeting a gene were pooled within each comparison. Severity was defined as median Log2 FC, whereas sensitivity and EK₅₀ were estimated from parent full-match and mismatch-guide series using predicted relative efficacies, with full-match guides assigned efficacy 1. Sensitivity was the predicted guide efficacy at which binned median fitness effects showed a sustained decrease to log₂FC ≤ −1. EK₅₀ was estimated from log-logistic fits as the predicted efficacy producing half the fitted fitness effect at efficacy 1. Emax was the fitted log₂FC at predicted guide efficacy 1, and EK₅₀ was the predicted guide efficacy producing half this effect relative to log₂FC = 0. Untreated consensus severity, sensitivity and EK₅₀ were calculated as medians of valid estimates across the four T1 batches (B1–B4; Supplementary Table S5). Consensus EK₅₀ was reported only when valid estimates were available in at least two batches.

### Guide-efficacy prediction model

A lasso model was trained on 6,868 full-match guides from 460 fitness affecting genes (having at least 5 full-match guides) using within-gene-centred depletion, with predictors encoding the 20-nt spacer, the variable N of the NGG PAM, 15 nucleotides downstream of the PAM and log-transformed target offset. Performance was assessed using 100 repeated 80:20 gene-held-out splits, with cross-validation for penalty selection within each training set. Illustrative validation panels used one representative split, while comparisons with CRISPRi-Pasteur and CIAO (MERF-based predictor) used common scored guides and repeated held-out predictions. Candidate guides were prioritized sequentially by target-position band, the *P. aeruginosa*-trained score, off-target burden and finally other guide-efficacy prediction models ranks, retaining up to 15 guides per gene.

## SUPPLEMENTARY FIGURES

**Supplementary Figure S1.**
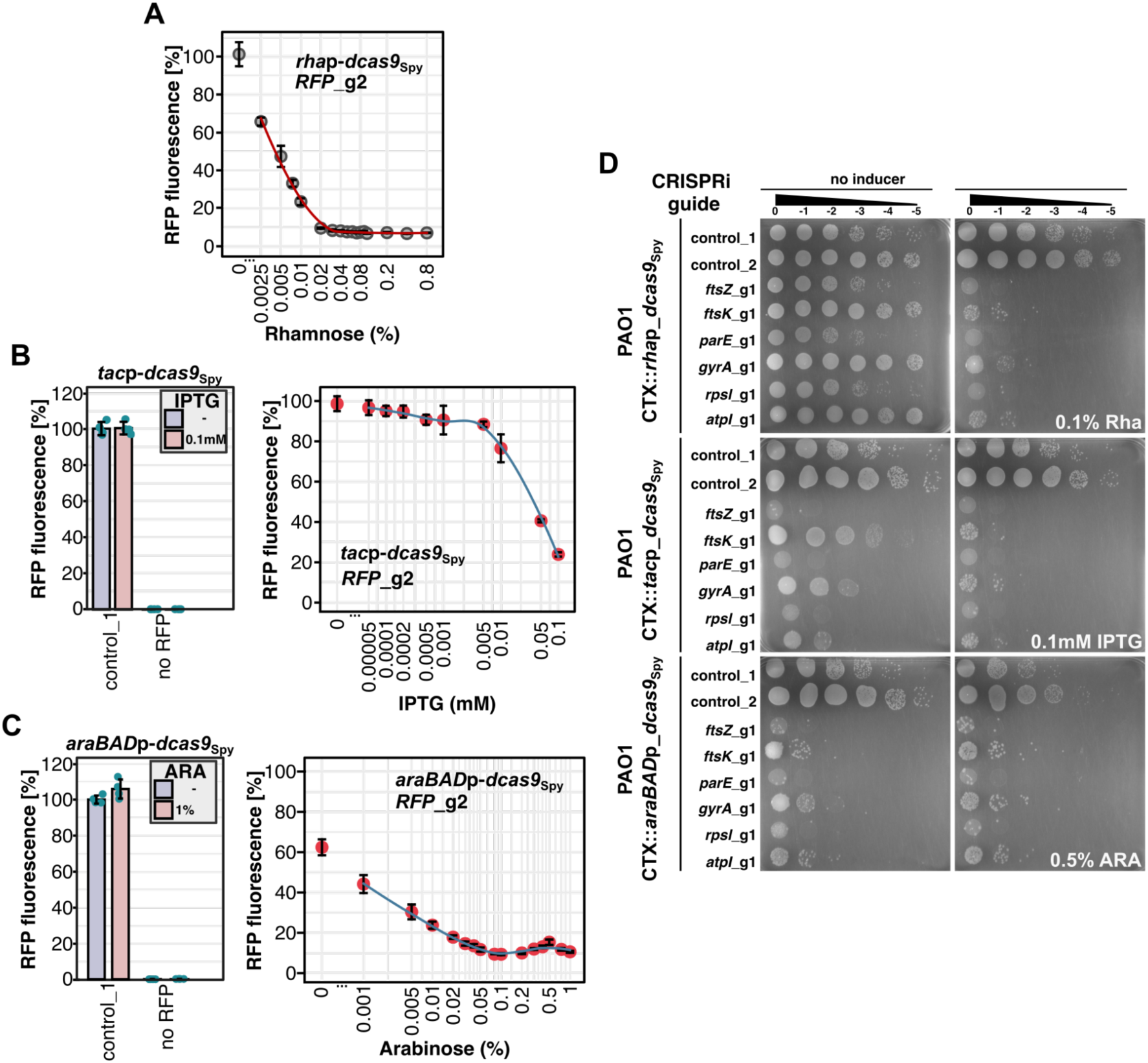
Comparison of inducible promoters for regulated dCas9*_Spy_*-dependent CRISPRi in *P. aeruginosa* PAO1. **(A–C)** Reporter-based evaluation of the rhamnose-inducible *rha*p (**A**), IPTG-inducible *tacp* (**B**) and arabinose-inducible *araBAD*p (**C**) for control of dCas9_Spy_ production. PAO1 strains carrying a constitutively expressed mRFP reporter, the corresponding promoter–*dcas9*_Spy_ cassette, and either a non-targeting or mRFP-targeting guide (*RFP*_g2) were grown with or without the corresponding inducer. mRFP fluorescence was normalized to OD₆₀₀ and expressed relative to the uninduced non-targeting control_1 strain, set to 100%. Data represent mean ± SD from four biological replicates. For **(A)** the control bar plots are the same as on Fig. 1B. **(D)** Comparison of basal and induced CRISPRi activity during essential-gene targeting. The indicated non-targeting or essential-gene targeting sgRNA vectors were introduced into PAO1 strains expressing dCas9_Spy_ from *rha*p, *tac*p or *araBAD*p. Conjugation mixtures were serially diluted and spotted on LB agar containing gentamicin and triclosan, with or without the corresponding inducer.

**Supplementary Figure S2.**
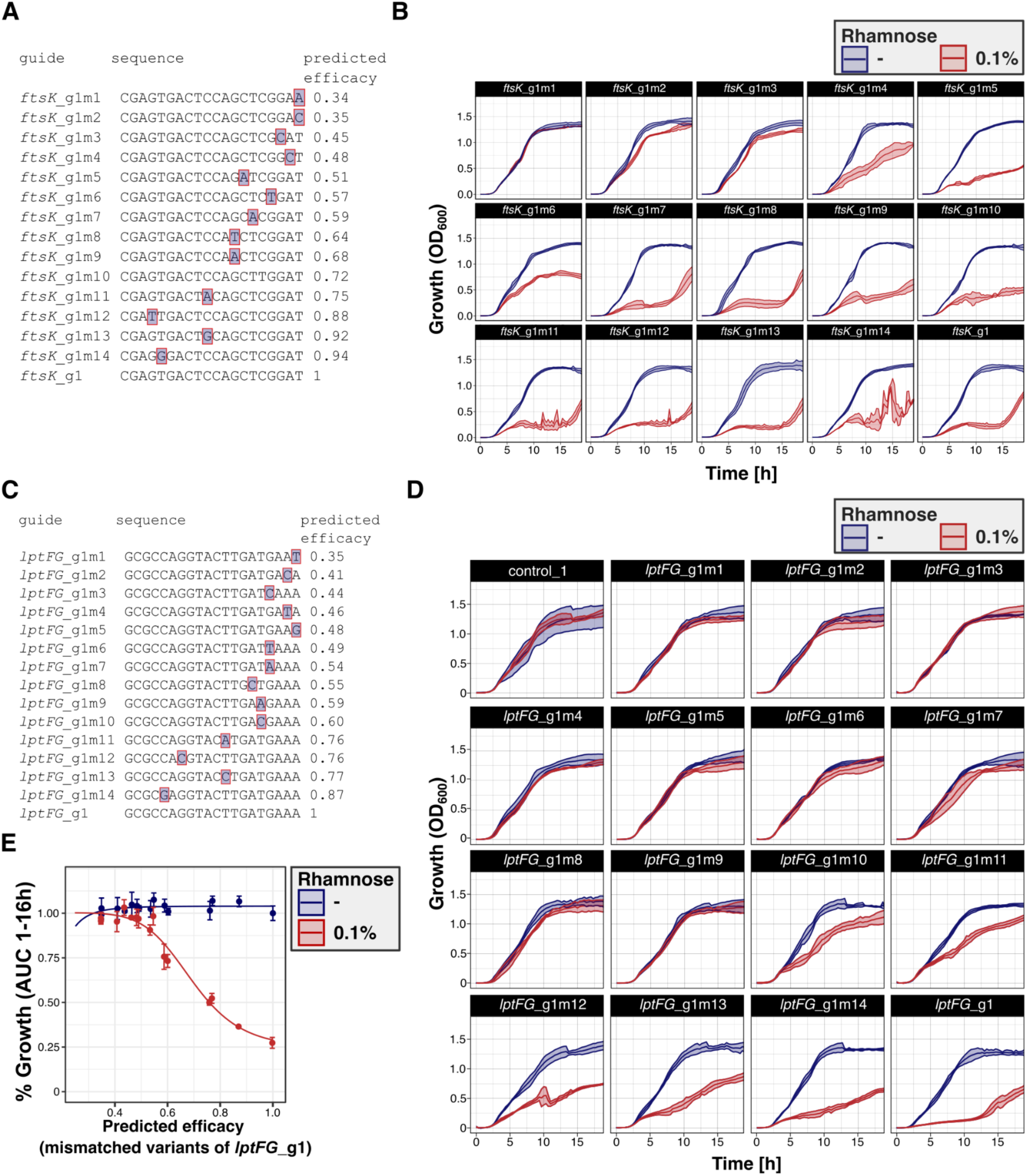
Mismatch-based modulation of essential-gene repression and its effects on PAO1 growth. **(A–B)** Mismatch-based modulation of *ftsK* repression. (**A**) Design of the *ftsK*_g1 mismatch series, comprising the full-match guide and single-nucleotide mismatch variants spanning a range of predicted silencing efficiencies. Boxes indicate the positions and identities of the introduced mismatches. (**B**) Growth of PAO1 strains carrying the indicated guides in MH medium with or without rhamnose. Curves represent the mean OD₆₀₀ ± SD from four biological replicates. The corresponding area-under-the-curve values are summarized in Fig. 1G. **(C–E)** Mismatch-based modulation of repression of the *lptF*-*lptG* operon. (**C**) Design of the *lptFG*_g1 mismatch series. (**D**) Growth of PAO1 strains carrying the full-match guide or the indicated mismatch variants in MH medium with or without rhamnose. (**E**) Relationship between predicted guide efficiency and the resulting growth effect, summarized using the area under the growth curve. Data represent the mean ± SD from up to four biological replicates.

**Supplementary Figure S3.**
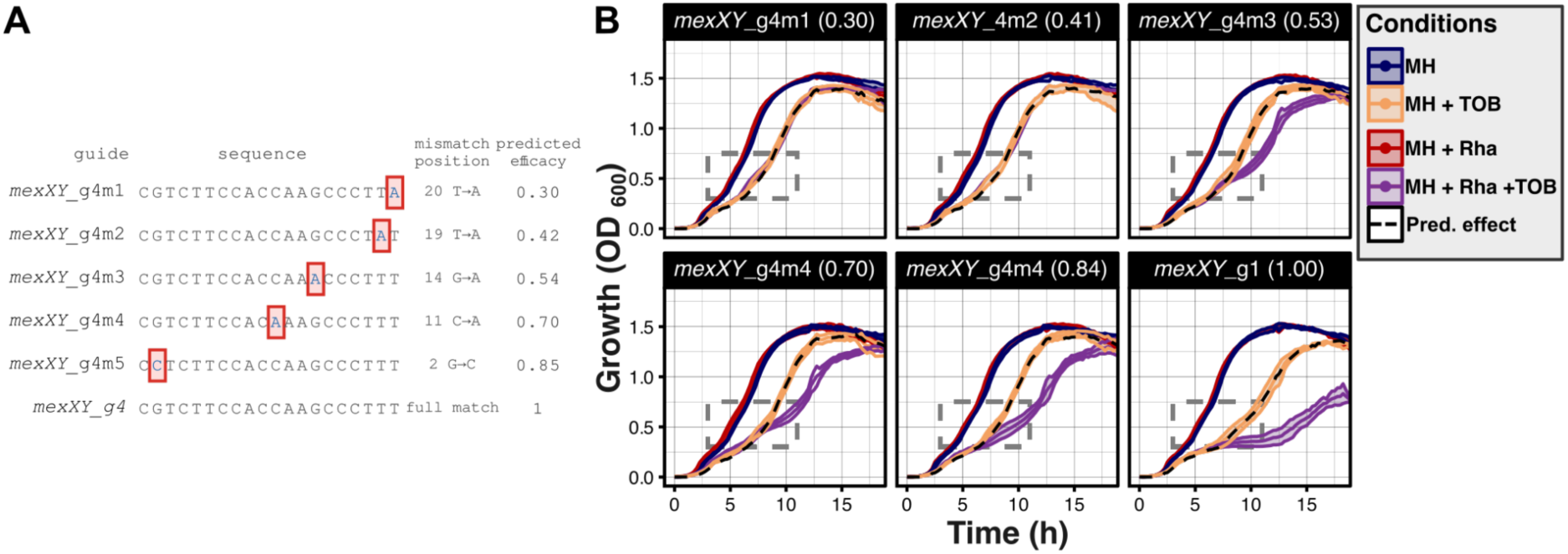
Mismatch-based modulation of *mexXY* repression and its effect on strain growth in the presence of tobramycin. **(A)** Design of the *mexXY*_g4 mismatched guide series, comprising the full-match guide and single-nucleotide mismatch variants. Red boxes indicate the positions and identities of the introduced mismatches. **(B)** Growth of PAO1 strains carrying the full-match *mexXY*_g4 guide or the indicated mismatch variants in MH medium containing subinhibitory 0.4 μg/mL TOB, with or without CRISPRi inducer (0.1% rhamnose; Rha). Curves represent mean OD₆₀₀ ± SD from up to fourbiological replicates. The dashed rectangle indicates the OD_600_/time interval used to calculate growth rates presented in Fig. 1H.

**Supplementary Figure S4.**
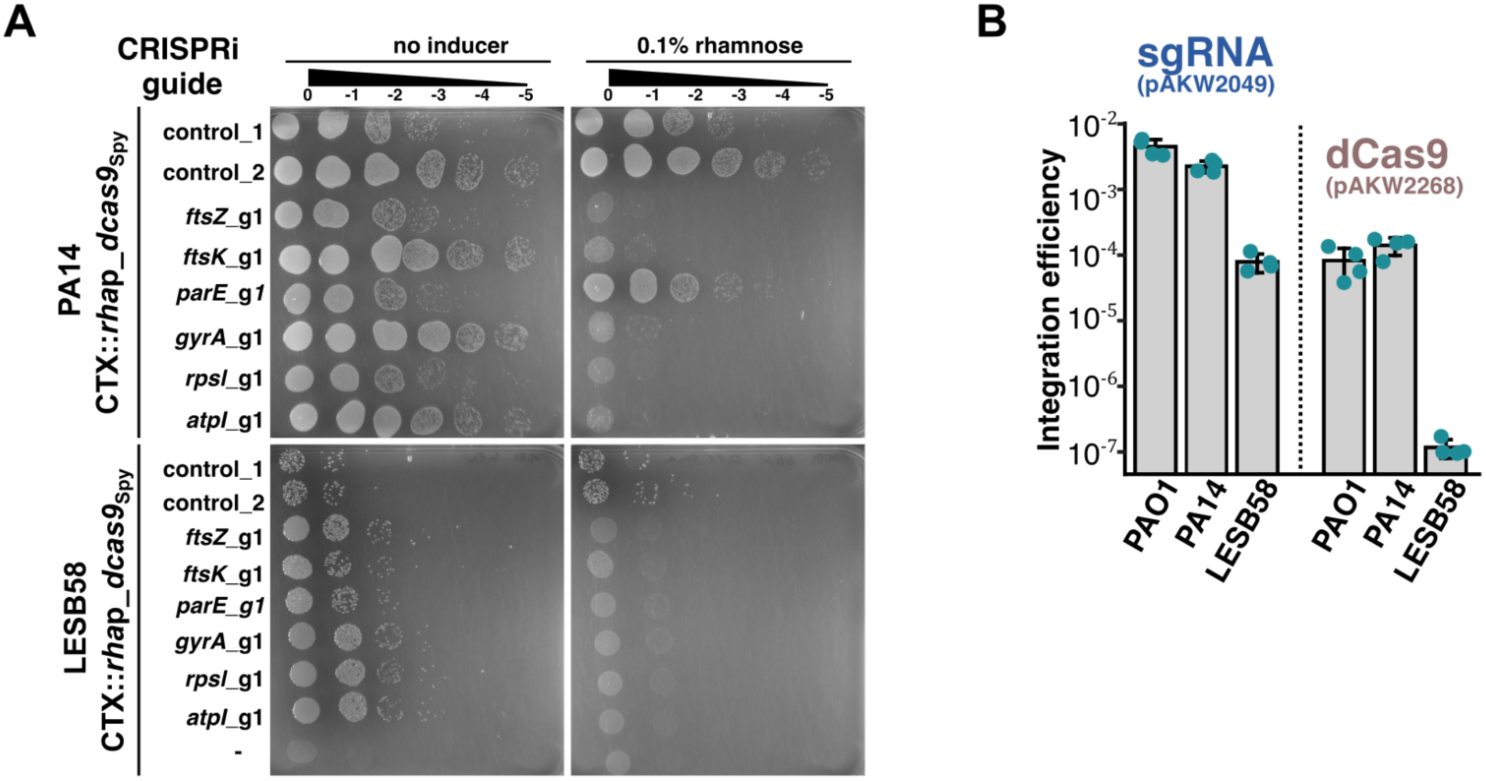
Vector integration and CRISPRi-mediated essential-gene repression across *P. aeruginosa* strains. **(A)** Essential-gene targeting in PA14 and LESB58. The indicated non-targeting or essential-gene-targeting sgRNA vectors were introduced into strains carrying the chromosomal *rha*p–*dcas9*_Spy_ cassette. Conjugation mixtures were spotted on selective LB agar with or without rhamnose to assess induction-dependent growth inhibition. Note that *parE*_g1 is not functional in PA14 because the selected sequence contains two mismatches in the extreme PAM-proximal nucleotides within the seed region. (-) indicates recipient and helper strain mixture. **(B)** Integration efficiency of the *dcas9*_Spy_-harbouring mini-CTX and sgRNA-cassette harbouring mini-Tn7 vectors in PAO1, PA14 and LESB58. Integration efficiency was calculated as the number of antibiotic-resistant integrants relative to the number of recipient cells. PAO1 and PA14 recipient cultures were grown at 42°C before conjugation, whereas LESB58 was grown at 37°C. Data represent mean ± SD from four biological replicates.

**Supplementary Figure S5.**
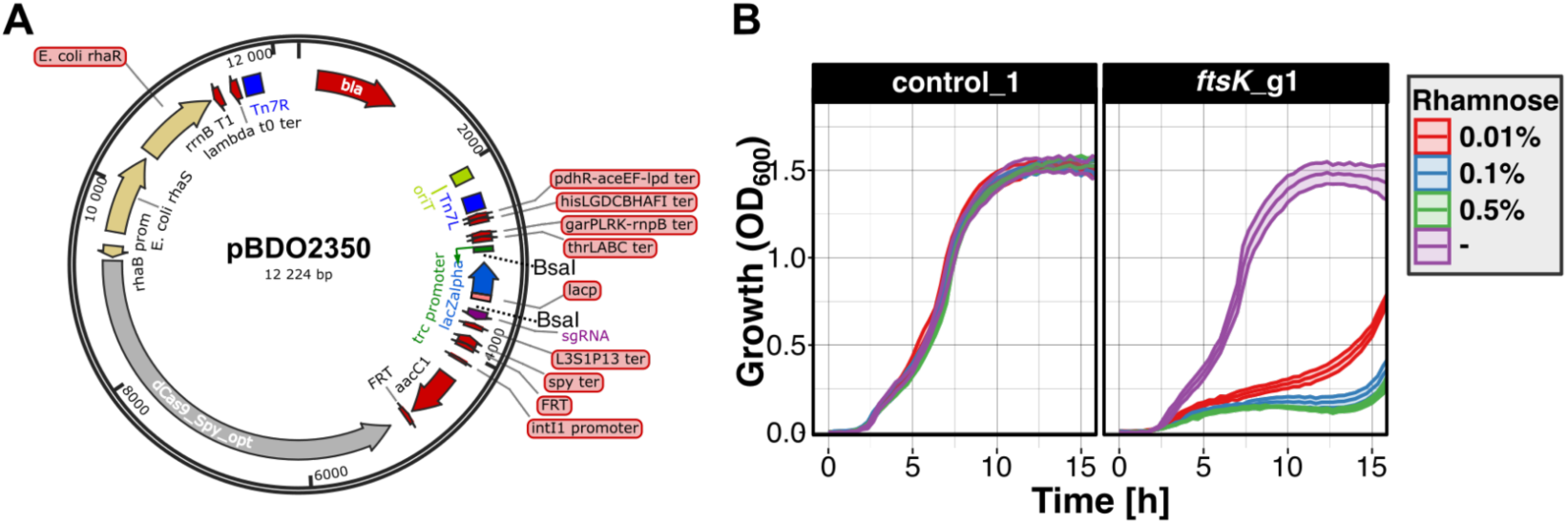
Single Tn7 based CRISPRi vector for *P. aeruginosa*. **(A)** Schematic representation of the mini-Tn7–based CRISPRi vector carrying both the rhamnose-inducible *rha*p-*dcas9_Spy_*cassette and the sgRNA-expression cassette. Spacer insertion by BsaI-mediated Golden Gate assembly replaces a *lacZα* cassette, facilitating identification of spacer-containing clones. **(B)** Functional validation of the single-vector CRISPRi system in PAO1. Strains carrying the control_1 or *ftsK*_g1 guide were grown with or without rhamnose. Growth curves represent OD₆₀₀ over time and data represent mean ± SD from *4* biological replicates.

**Supplementary Figure S6.**
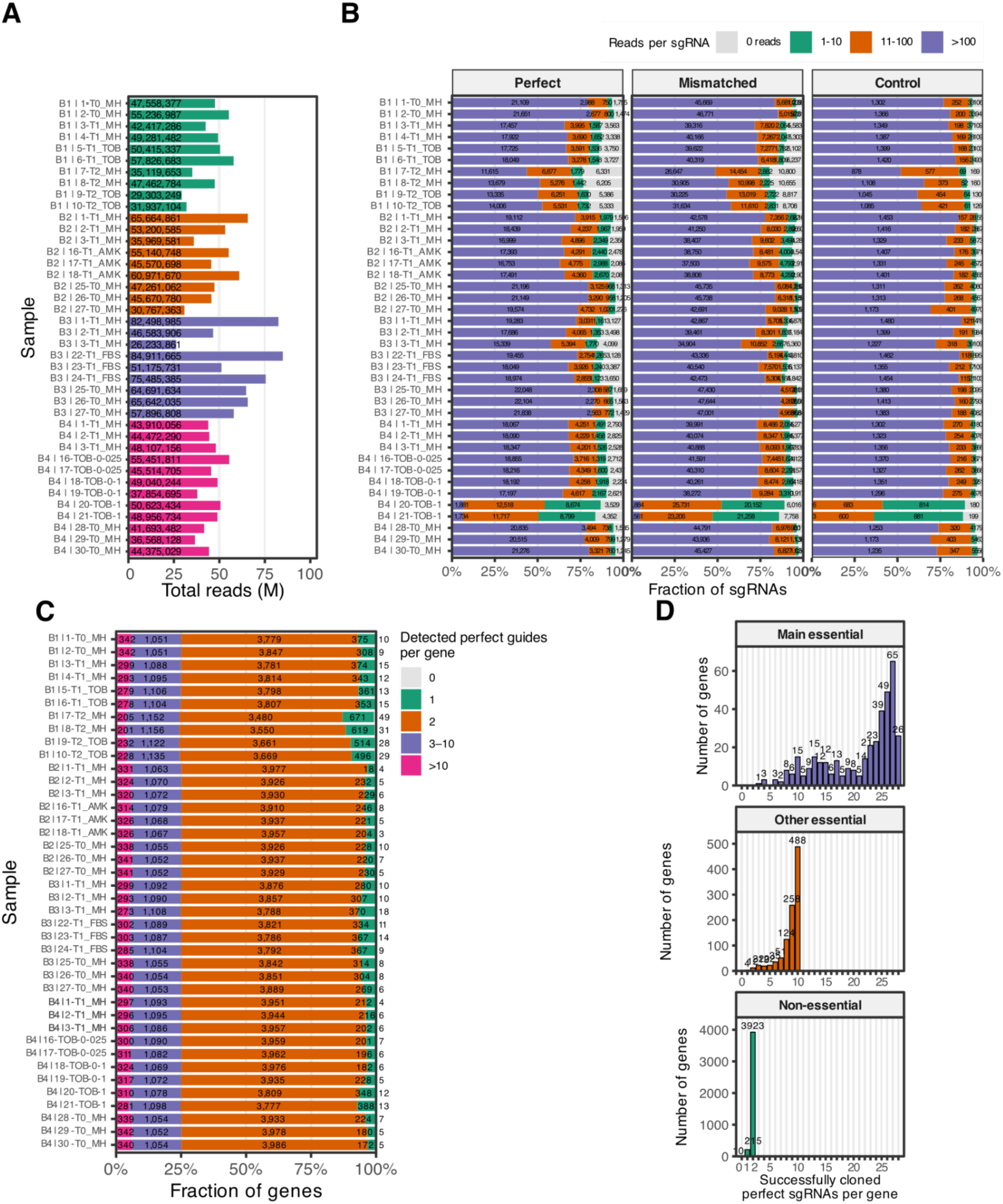
Sequencing coverage and cloning success of the CRISPRi library. (**A**) Total assigned sgRNA reads across samples included in the CRISPRi-seq experiments. B1-B4 indicate independent batches. (**B**) Distribution of sgRNA read depth in each sample, shown separately for full-match, mismatched and non-targeting control sgRNAs. Guides were grouped into 0, 1–10, 11–100 and >100-read categories. (**C**) Number of detected full-match sgRNAs per gene in each sample. A guide was considered detected when represented by at least one read. (**D**) Distribution of successfully cloned full-match sgRNAs per gene across the library-design groups. A guide was considered successfully cloned when its mean abundance in the no-drug T0 samples was at least 10 reads in at least one experimental batch. Category assignment is described in Supplementary Table S2.

**Supplementary Figure S7.**
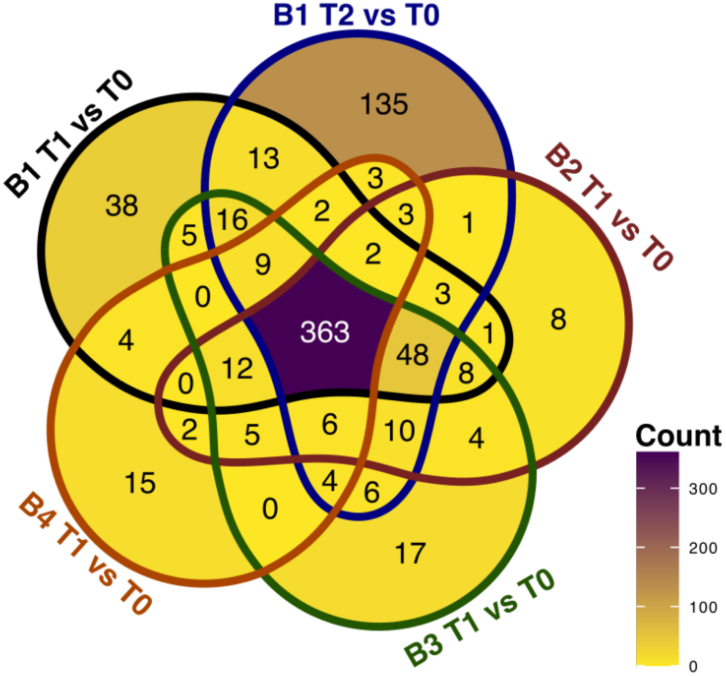
Overlap of CRISPRi-associated fitness defects in MH. Venn diagram showing genes with significant negative fitness effects across batches and time points in induced, untreated MH cultures, relative to the corresponding T0 reference. Genes were included at Stouffer *p*< 0.05 and median log₂FC ≤ −1.

**Supplementary Figure S8.**
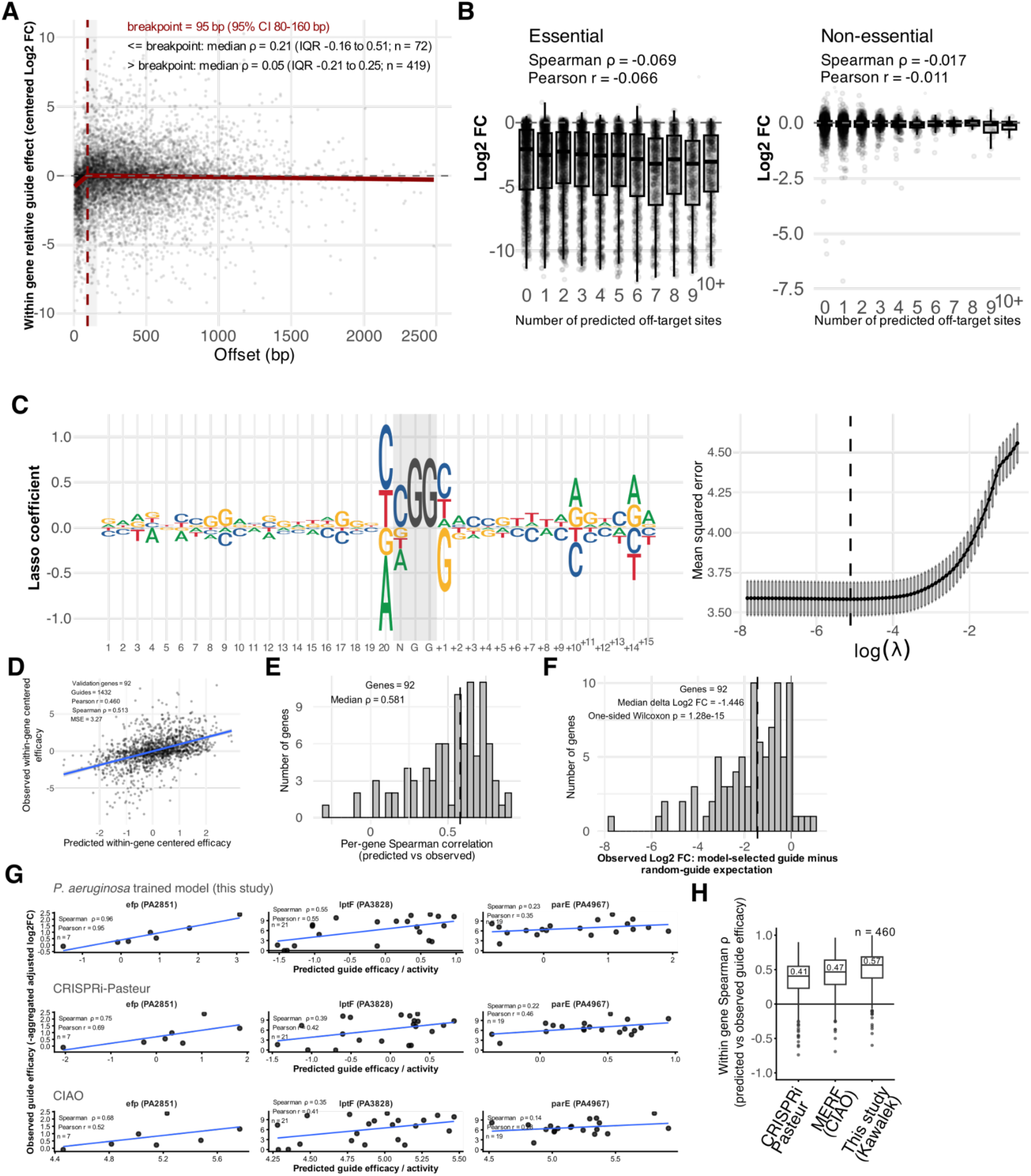
Determinants of CRISPRi guide efficacy. **(A)** Relationship between guide position and relative CRISPRi guide effect for full-match sgRNAs targeting genes showing a significant negative fitness effect in the untreated CRISPRi-seq comparisons. Guide effect was centred relative to other guides targeting the same gene, with more negative values indicating stronger relative effects. The breakpoint in the relationship between offset and guide effect was estimated by continuous piecewise regression. For each gene, Spearman correlations between distance from the start codon and relative guide effect were calculated separately for guides targeting sites before and after the breakpoint. **(B)** Effect of predicted off-targeting on CRISPRi guide performance. Off-target number was defined as the number of genomic sites with perfect complementarity to the nine PAM-proximal nucleotides of the sgRNA spacer, as described previously (41). Boxplots show the distribution of aggregated adjusted log₂ fold changes for full-match sgRNAs according to predicted off-target count, shown separately for genes with negative fitness effects (Supplementary Table S5) and the remaining genes. Boxes indicate the median and interquartile range, with individual guides shown as points. **(C)** Lasso regression model of CRISPRi guide efficacy. Sequence-feature coefficients summarize the relative contribution of nucleotides at individual positions within the spacer, the variable N position of the NGG PAM, and the downstream sequence context, with larger positive coefficients indicating increased predicted guide efficacy. The panel on the right represents cross-validation curve - mean squared error as a function of the model regularization parameter (λ), with the selected value indicated. **(D)** Validation of predicted guide efficacy using genes excluded from model training. Predicted guide scores are plotted against observed fitness effects for individual full-match sgRNAs, with both expressed relative to other guides targeting the same gene. Mean squared error (MSE), Pearson, and Spearman correlation coefficients used for relationship analysis. **(E)** Distribution of within-gene Spearman correlations between predicted and observed guide efficacy across genes in the validation set. **(F)** Top-guide enrichment analysis in the validation set. The distribution shows the difference in observed adjusted log₂ fold change between the highest-ranked guide selected by the model and the random-guide expectation for each gene. Negative values indicate stronger fitness effects for model-selected guides. **(G)** Comparison of predicted guide activity with experimentally observed efficacy for the *P. aeruginosa*-trained model, CRISPRi-Pasteur and CIAO across three selected loci: *lptF*, *efp* (*PA2851*) and *parE* (*PA4967*). Each point represents one full-match sgRNA. Observed efficacy was defined as the negative aggregated adjusted log₂ fold change, such that higher values indicate stronger experimental effects. Model scores were oriented so that higher values consistently indicate stronger predicted activity. Spearman ρ and Pearson coefficients were calculated for each gene–model combination. **(H)** Distribution of guide-ranking performance across the *P. aeruginosa*-trained model, CRISPRi-Pasteur and CIAO. Analysis included 460 genes represented by at least five full-match sgRNAs with predictions available from all three models. Boxplots show the distribution of within-gene Spearman correlations between predicted and experimentally observed guide efficacy. Boxes indicate the median and interquartile range, whiskers extend to 1.5 × the interquartile range, and points indicate outliers. All three models were compared using the same guides. For the *P. aeruginosa*-trained model, predictions were combined across 100 repeated splits into training and test sets, using only predictions for genes excluded from both training and parameter selection.

**Supplementary Figure S9.**
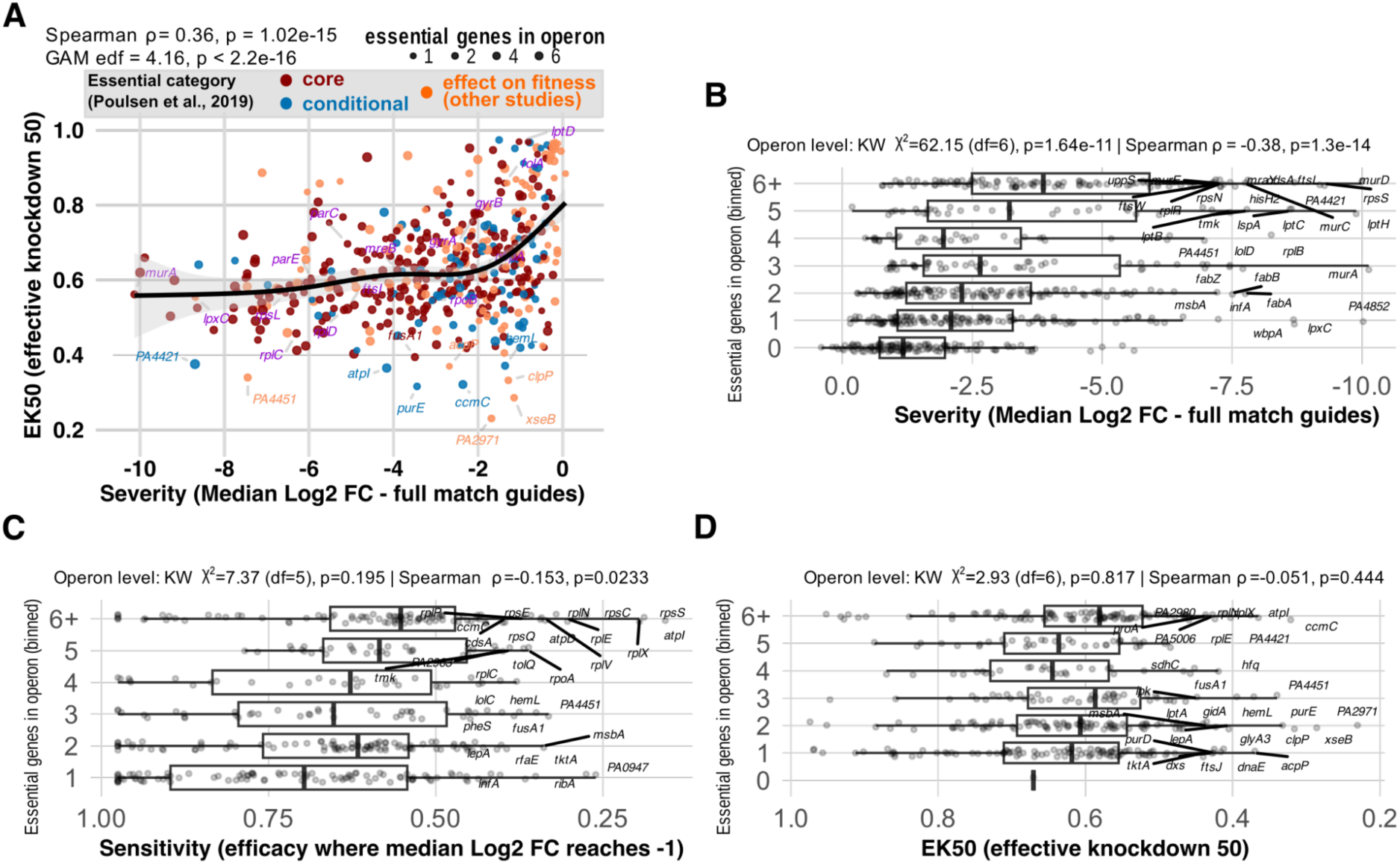
Relationships among CRISPRi gene-vulnerability measures and their association with transcriptional-unit essential-gene content. **(A)** Relationship between severity and EK₅₀ estimated from untreated T1-versus-T0 comparisons. Severity was calculated as the median adjusted log₂ fold change of full-match sgRNAs, whereas EK_50_ (effective knockdown 50) represents the predicted knockdown required to produce half of the maximal fitted fitness effect. Values represent consensus estimates across the four T1 vs T0 untreated library comparisons (Supplementary Table S5), and EK_50_ was reported only when valid estimates were obtained in at least two comparisons. Only genes with a numeric consensus EK_50_ are shown. Points are colored according to essentiality category and scaled according to the number of essential genes within the corresponding transcriptional unit. Selected antimicrobial targets and genes with extreme values are labelled. The black line and shading indicate a generalized additive model fit and 95% confidence interval. Spearman ρ indicates the strength and direction of the monotonic association, with the corresponding *p*-value testing whether this association differs from zero. The GAM effective degrees of freedom (edf) is used as a descriptor of the shape of the fitted relationship. **(B)** Distribution of CRISPRi severity according to the number of essential genes within the annotated transcriptional unit. Analysis was restricted to genes showing a significant negative fitness effect in at least one untreated T1-versus-T0 comparison. Transcriptional units were grouped according to the number of essential genes they contained (based on the previous Tn-seq based analyses - Supplementary Table S2), with those containing six or more essential genes combined into a single category. Kruskal–Wallis tests and Spearman correlations used one summary value per transcriptional unit. Statistical tests used the median severity, sensitivity or EK₅₀ of the included genes within each transcriptional unit. Individual gene values are shown for visualization Both analyses were performed using individual genes and repeated using one summary value per transcriptional unit, avoiding treatment of genes within the same unit as independent observations. **(C)** Relationship between sensitivity to partial repression and transcriptional-unit essential-gene content for genes with an estimated sensitivity. Data representation and analyses are as in **(B)**. **(D)** Relationship between EK50 and TU essential-gene content for genes with a numeric EK50 estimate. Data representation and statistical analyses are as in panel **(B)**.

**Supplementary Figure S10.**
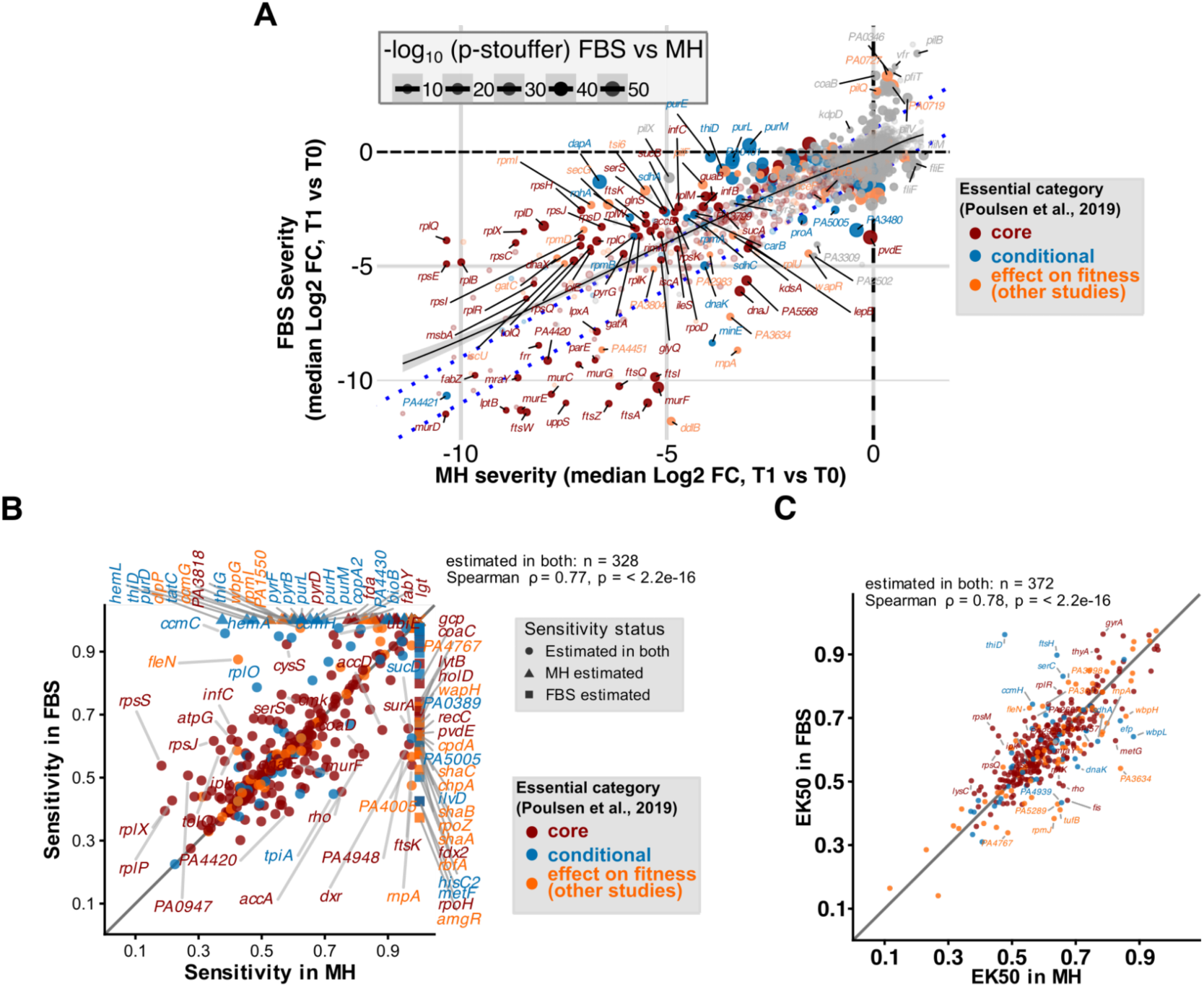
Comparison of CRISPRi gene-vulnerability measures between MH and FBS. **(A)** Comparison of gene-level CRISPRi severity in MH and FBS. Each point represents one gene positioned according to its median log₂ fold change relative to T0 in MH and FBS. Point color indicates essentiality category. Genes showing significant condition-dependent effects are labelled. The diagonal indicates equal effects between conditions, and dashed lines indicate ±1 log₂FC differences. **(B)** Comparison of sensitivity to partial repression in MH and FBS for genes showing a significant negative fitness effect relative to T0 in at least one of the two conditions. Sensitivity was estimated independently in each condition as the predicted sgRNA efficacy at which the median fitness defect showed a sustained decrease to log₂FC ≤ −1. Genes that were evaluable but did not reach the sensitivity threshold are displayed at the upper boundary of 1 and distinguished by point shape. Genes lacking sufficient guide or efficacy-bin information are excluded. Point color indicates essentiality category. Correlation and statistical analyses include only genes with numeric sensitivity estimates in both conditions. **(C)** Comparison of EK₅₀ between MH and FBS for genes showing a significant negative fitness effect relative to T0 in both conditions. EK₅₀ was estimated independently in each condition from fitted relationships between predicted sgRNA efficacy and fitness effect. Only genes with valid estimates in both conditions are shown (*n* = 332). Point colour indicates essentiality category, and the diagonal line indicates equal EK₅₀ values. Spearman correlation and a paired Wilcoxon signed-rank test assessed the association and differences between conditions, respectively.

**Supplementary Figure S11.**
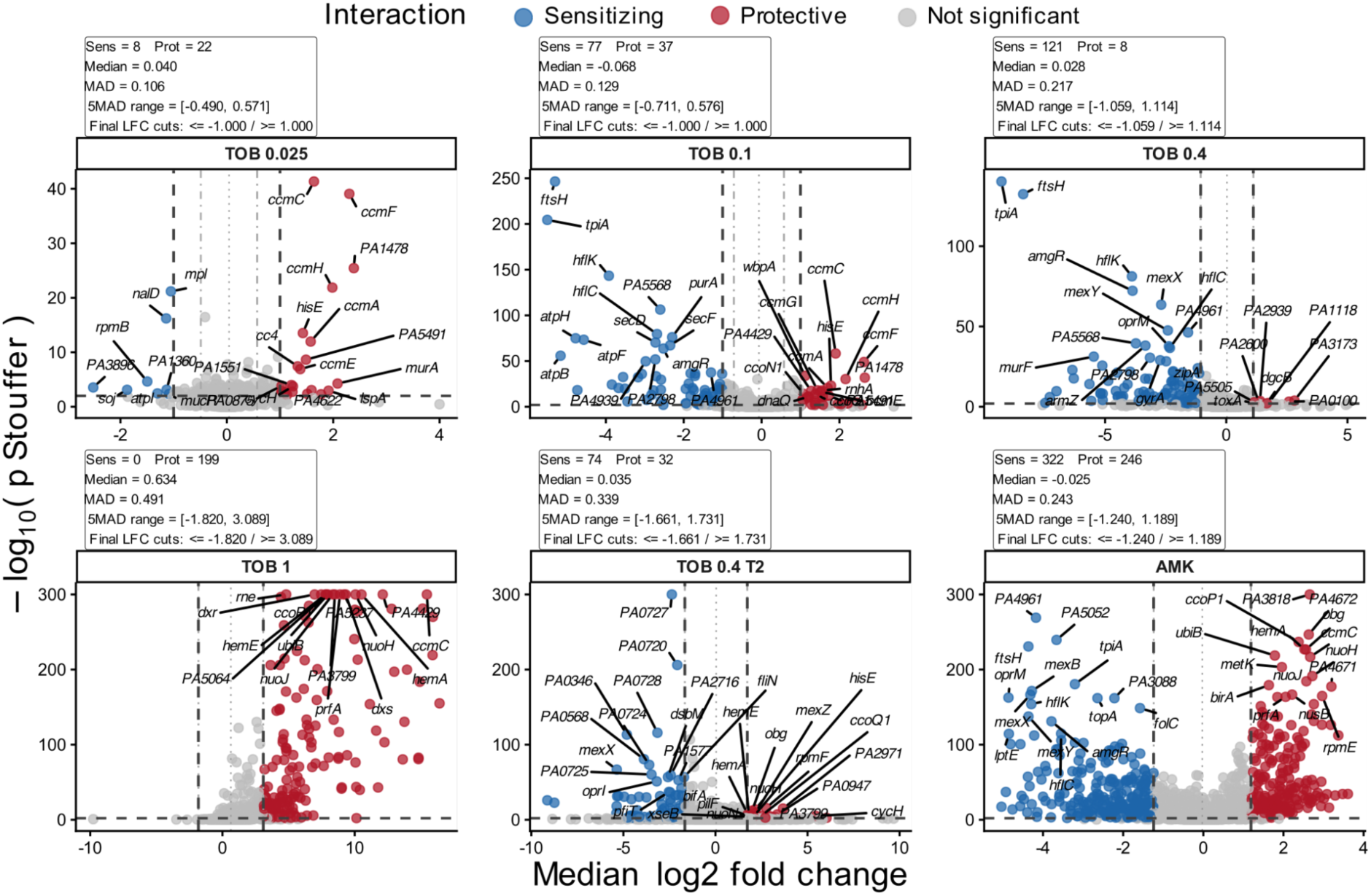
Gene–drug interactions identified across different antibiotic treatments. Volcano plots display gene-level changes in response to treatment with various concentrations of tobramycin (TOB) and amikacin (AMK). The x-axis shows interaction severity (median log2 fold change), and the y-axis shows −log10 of the Stouffer-combined *p* value. The horizontal dashed line marks Stouffer *p* < 0.01, while vertical dashed lines indicate condition-specific log2 fold change cutoffs defined as the median ± 5 median absolute deviation (MAD), with a minimum absolute cutoff of 1. Genes meeting both the *p* value and direction-specific log2 fold change threshold were classified as sensitizing or protective. Up to 15 genes are labelled per condition, prioritizing lower *p* values and larger absolute log2 fold change values. Tables above the plots indicate statistics of identified interactions and exact cutoff values used.

**Supplementary Figure S12.**
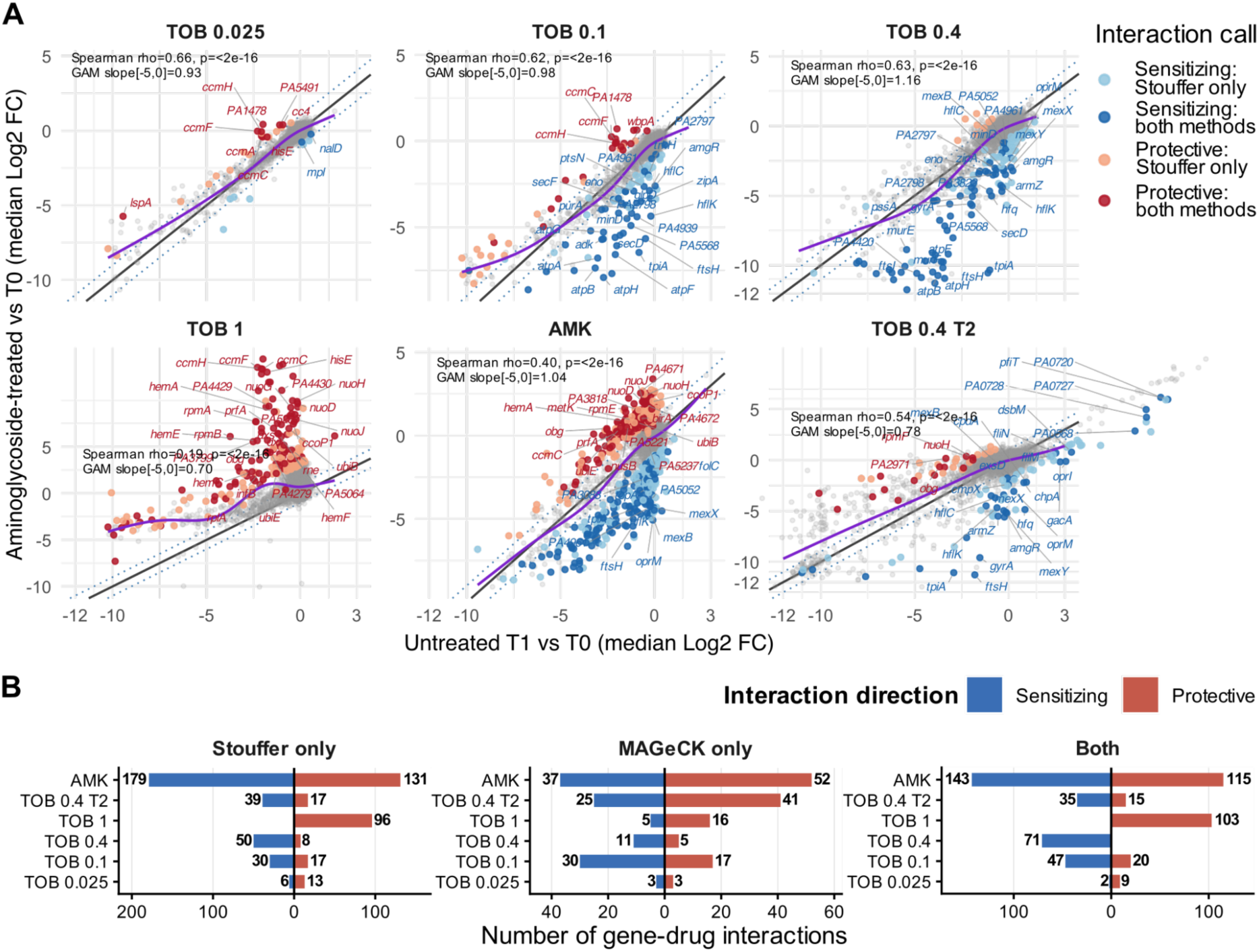
Relationship between untreated and aminoglycoside-treated CRISPRi repression-triggered fitness effects and comparison of gene–drug interaction calling methods. **(A)** Gene-level fitness effects in matched untreated cultures and under aminoglycoside treatment, each relative to T0. For each condition, the median log₂ fold change in the matched untreated control is plotted on the x-axis against the corresponding value under aminoglycoside treatment on the y-axis. Each point represents one gene. Significant gene–drug interactions were identified using Stouffer *p* < 0.01 and condition-specific medLFC thresholds defined by median ± 5 MAD, with a minimum absolute cutoff of 1. Interactions independently identified in the same direction by MAGeCK RRA (FDR < 0.05) are shown in darker colors, whereas Stouffer-only interactions are shown in lighter colors and nonsignificant genes in grey. Gene labels are restricted to interactions supported in the same direction by both methods. The solid diagonal indicates equal fitness effects in treated and untreated conditions, and dotted lines indicate ±1 log₂FC deviation. The purple line shows the GAM fit. Spearman correlation and the mean GAM slope over the untreated log₂ FC range −5 to 0 are reported for each condition. MAGeCK results are supplied in Supplementary Table S8. **(B)** Numbers of sensitizing and protective gene–drug interactions identified by the Stouffer-based analysis only, MAGeCK RRA only, or by both methods in the same direction, in each comparison. Bars indicate the number of genes in each category.

**Supplementary Figure S13.**
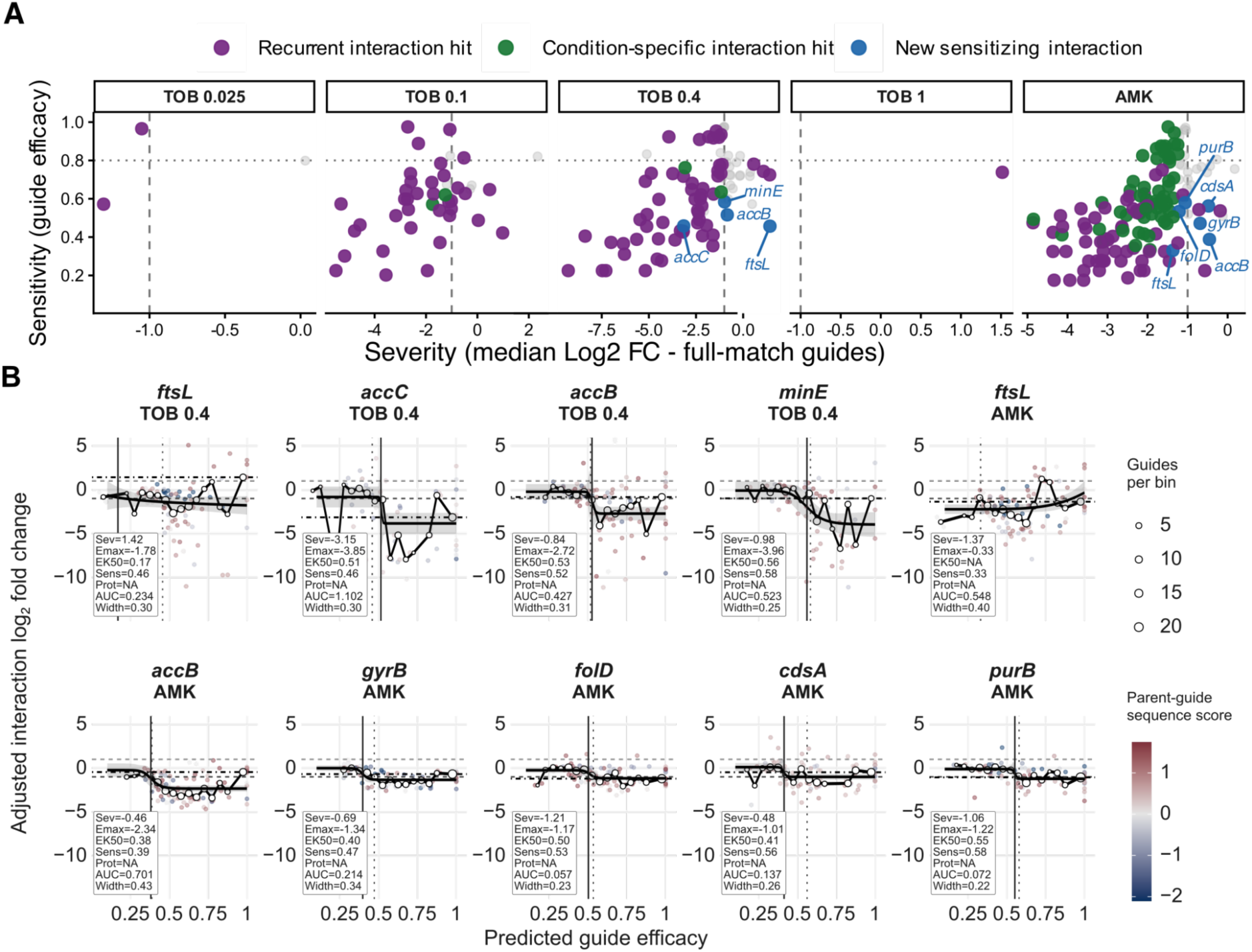
Mismatch-guide analysis of gene–aminoglycoside interactions. **(A)** Relationship between interaction severity and sensitivity for additional candidate sensitizing gene–drug interactions across aminoglycoside conditions. Candidates were required to reach the direction-specific interaction threshold at a predicted guide efficacy ≤0.8, lack a same-direction full-match gene-level call, be absent from the corresponding recurrent set, and show a sustained LOESS-fitted response beyond the threshold. **(B)** Relationship between predicted guide efficacy and adjusted interaction log₂ FC for the additional sensitizing candidates identified in panel (**A**). Sev - interaction severity, defined as the median log₂ FC of full-match guides. Emax - fitted interaction effect at maximal predicted guide efficacy. EK_50_ - predicted guide efficacy producing half of Emax. Sens and Prot, predicted-efficacy thresholds for sensitizing and protective interactions, respectively. AUC, area of the LOESS-fitted response beyond the interaction threshold. Width, predicted-efficacy range over which the LOESS fit remained beyond the threshold.

**Supplementary Figure S14.**
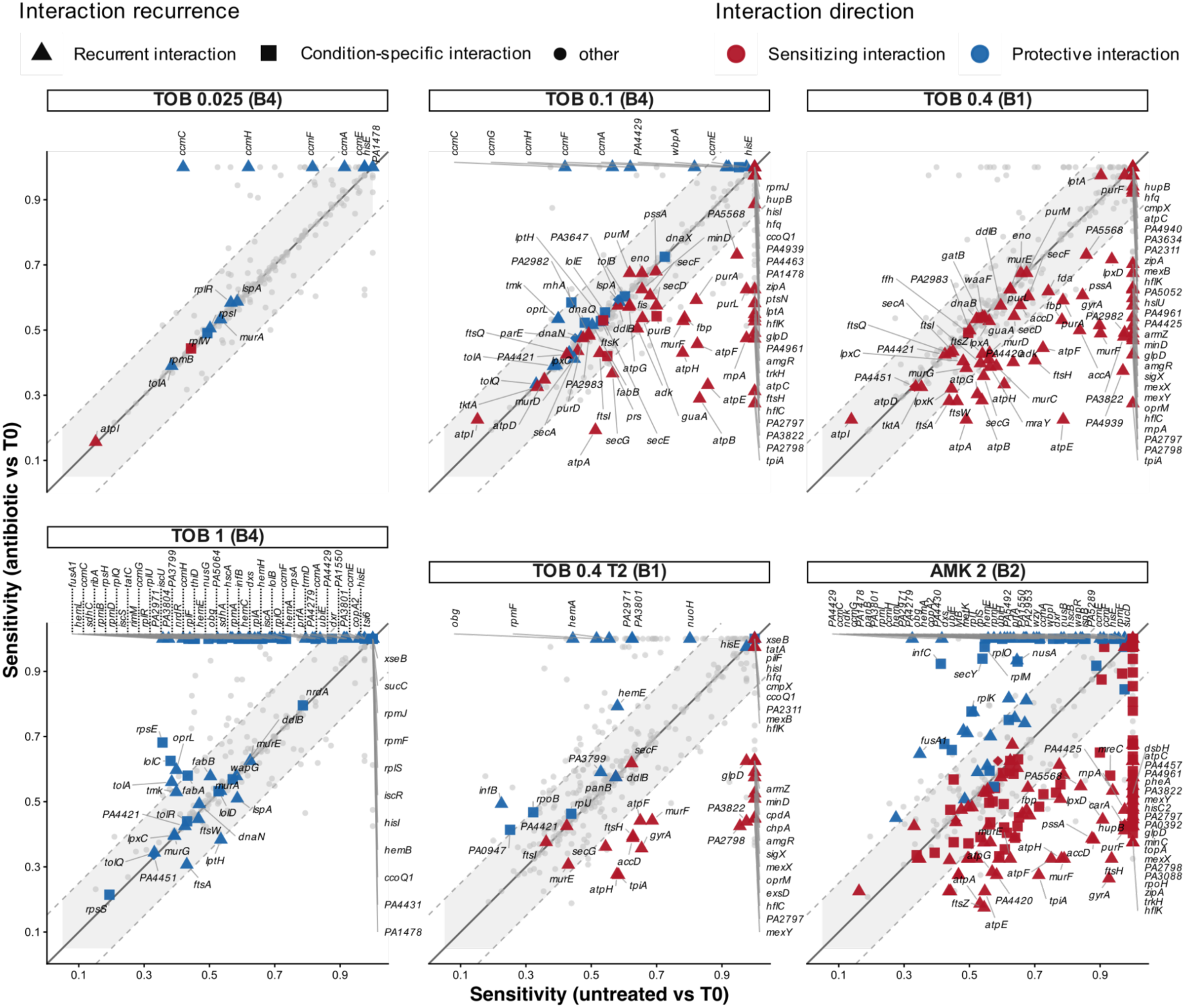
Aminoglycoside-associated shifts in gene repression thresholds. Predicted guide-efficiency thresholds were compared between matched untreated and antibiotic-treated conditions. Thresholds indicate the predicted level of gene silencing required to cause a sustained fitness defect (sensitivity; log₂ FC ≤ −1). Only genes with significant negative fitness effects in at least one matched condition and evaluable sensitivity profiles in both conditions were included. Grey points represent genes, whereas red and blue points indicate sensitizing and protective interactions, respectively (Supplementary Table S8). Point shapes distinguish recurrent interactions, condition-specific interactions, and non-recurrent interactions. The solid diagonal indicates equal thresholds, and dashed lines indicate differences of ±0.15. Genes without a threshold estimate at one comparison are displayed at 1. Significant interaction genes are labeled. for AMK, only those with largest threshold shifts are labeled.

**Supplementary Figure S15.**
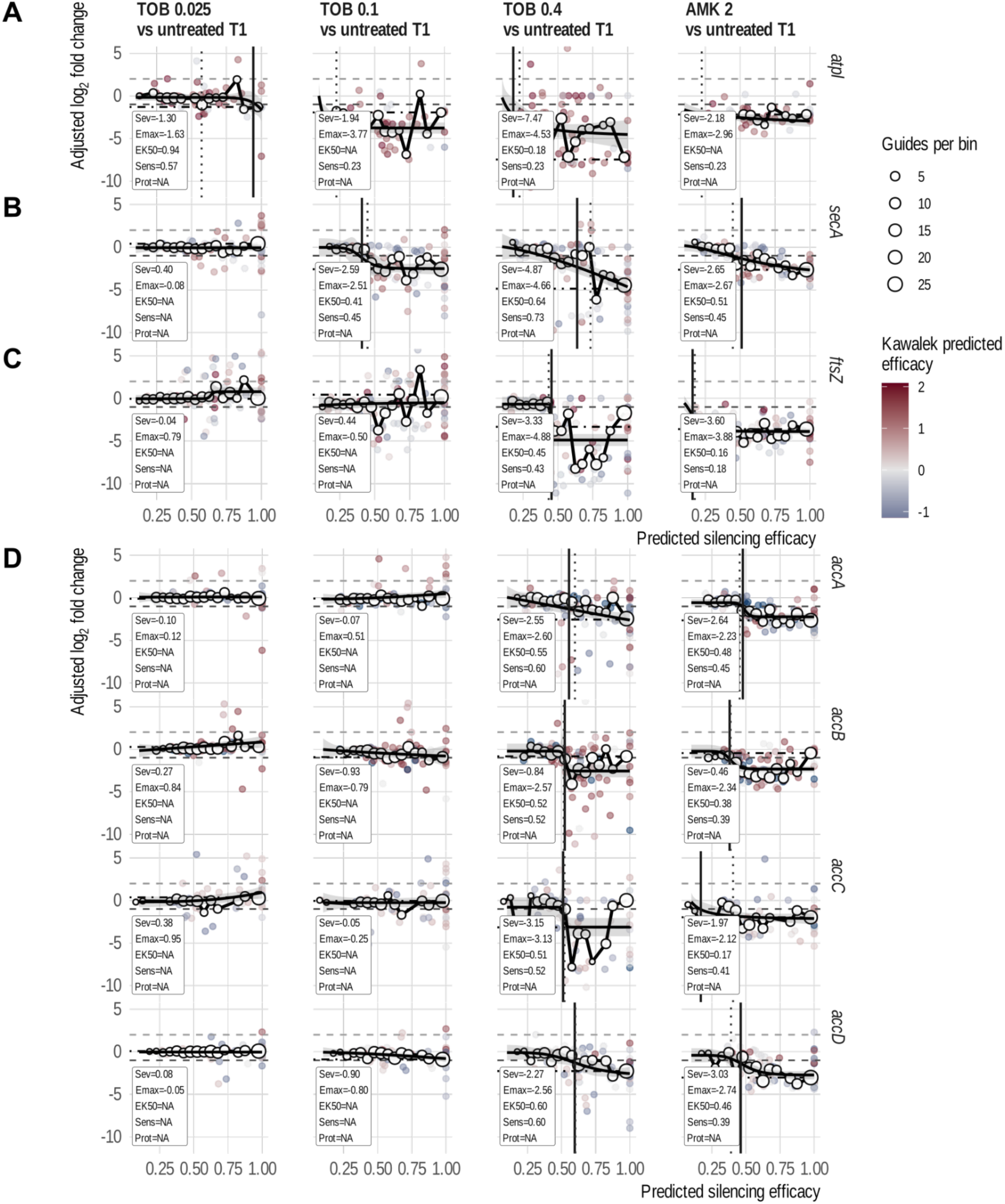
Guide-efficacy dependence of aminoglycoside interactions for selected genes, including those selected for experimental validation. **(A)** *atpI*, **(B)** *secA*, **(C)** *ftsZ*, and **(D)** Acetyl-CoA carboxylase genes *accA, accB, accC* and *accD*. Full-match and mismatch log₂ fold change of abundance was plotted relative to predicted silencing efficacy, across the indicated aminoglycoside T1 vs untreated T1 comparisons. Full-match guides were assigned an efficacy of 1. Binned median effects summarize the relationship between predicted silencing efficacy and the interaction phenotype, with horizontal dashed lines indicating the fitness change thresholds. Parameter Emax denotes the fitted adjusted log₂ fold-change at maximal predicted silencing efficacy, representing the estimated maximal interaction effect derived from the fitted curve. Sev, interaction severity (median log2 fold change of full-match guides). EK_50_, predicted guide efficacy producing half of Emax. Sens and Prot, sensitizing or protective thresholds, respectively.

**Supplementary Figure S16.**
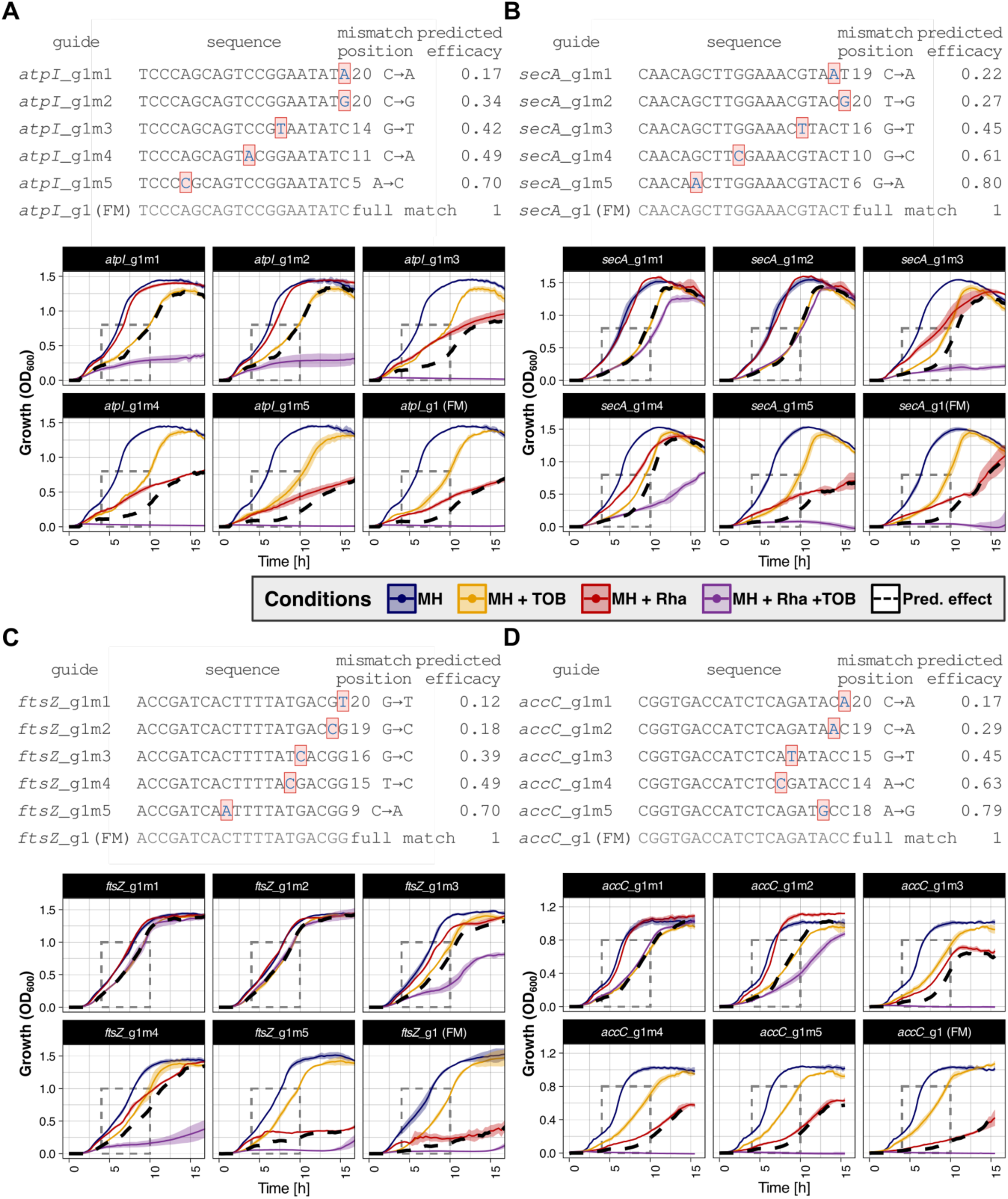
Mismatch-guide design and complete growth profiles used for validation of aminoglycoside. interactions**. (A–D)** Complete growth profiles for CRISPRi guide series targeting *atpI* (**A**), *secA* (**B**), *ftsZ* (**C**) and *accC* (**D**). For each gene, the top panel shows the full-match guide and five mismatch derivatives, including guide sequences, mismatch positions and predicted silencing efficacies. Substituted nucleotides are highlighted relative to the full-match guide. Strains carrying each guide were grown in MH, MH containing 0.4 µg/mL TOB, MH containing 0.1% rhamnose, or MH containing both rhamnose and TOB. Growth curves show mean OD_600_ ± SD across biological replicates. The dashed box indicates the predefined interval used for growth-rate calculation, and the black dashed curve represents the multiplicative expectation calculated from the corresponding single-treatment conditions.

**Supplementary Figure S17.**
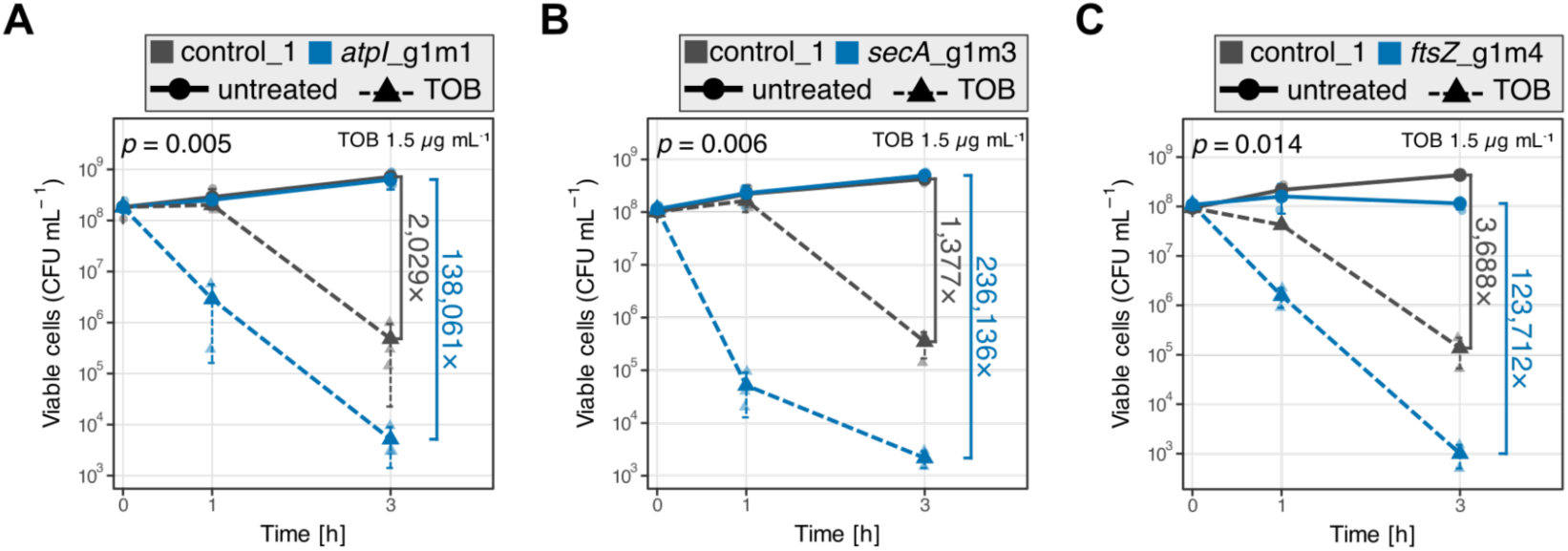
Validation of aminoglycoside sensitization in the PA14 genetic background. Tobramycin killing assays for PA14 strains carrying the corresponding *atpI*, *secA* and *ftsZ* mismatch guides used for validation in PAO1, shown in A–C, respectively. The *accC* construct was not tested in PA14 because the full-match guide selected in PAO1 contains a mismatch to the corresponding PA14 target sequence. Strains were precultured under CRISPRi-inducing conditions (MH + 0.1% Rha) and subsequently transferred to medium with or without TOB at 1.5 μg mL⁻¹. Viability was determined as CFU mL⁻¹ at the indicated time points. Data show mean ± SD from three biological replicates, with individual replicates displayed. 3-hour fold decreases represent the reduction in viable counts following TOB exposure. Replicate-specific log₁₀ CFU reductions were compared between each knockdown and the non-targeting control using a paired two-sided *t*-test.

